# Deep mutational scanning reveals sequence to function constraints for SWEET family transporters

**DOI:** 10.1101/2024.06.28.601307

**Authors:** Krishna K. Narayanan, Austin T. Weigle, Lingyun Xu, Xuenan Mi, Chen Zhang, Li-Qing Chen, Erik Procko, Diwakar Shukla

## Abstract

Protein science is entering a transformative phase enabled by deep mutational scans that provide an unbiased view of the residue level interactions that mediate function. However, it has yet to be extensively used to characterize the mutational and evolutionary landscapes of plant proteins. Here, we apply the method to explore sequence-function relationships within the sugar transporter AtSWEET13. DMS results describe how mutational interrogation throughout different regions of the protein affects AtSWEET13 abundance and transport function. Our results identify novel transport-enhancing mutations that are validated using the FRET sensor assays. Extending DMS results to phylogenetic analyses reveal the role of transmembrane helix 4 (TM4) which makes the SWEET family transporters distinct from prokaryotic SemiSWEETs. We show that transmembrane helix 4 is intolerant to motif swapping with other clade-specific SWEET TM4 compositions, despite accommodating single point-mutations towards aromatic and charged polar amino acids. We further show that the transfer learning approaches based on physics and ML based *In silico* variant prediction tools have limited utility for engineering plant proteins as they were unable to reproduce our experimental results. We conclude that DMS can produce datasets which, when combined with the right predictive computational frameworks, can direct plant engineering efforts through derivative phenotype selection and evolutionary insights.

## MAIN

Sugars Will Eventually be Exported Transporters (SWEETs) are bidirectional uniporters that transport both sugars and gibberellins for regulation of several plant developmental and physiological processes.^1–5^ Understanding the sequence-function space of SWEETs may inform new avenues of exploration for biotechnological applications, such as improved crop yield and reduced pathogen susceptibility.^3^ Aside from one study,^6^ the plant sciences field has yet to harness the power of deep mutational scanning as a generalizable methodology.^7^ Deep mutational scanning could prove critical to enhancing throughput of molecular-level discoveries in the plant sciences.^8^ Here, we perform a mutagenesis of SWEET13 from *Arabidopsis thaliana* (AtSWEET13) to identify sites of interest for enhanced cellular influx of the fluorescent sucrose analogue, esculin.^9,10^ The conserved residues correlate well with established structural and mechanistic studies of AtSWEET13, suggesting the scan was able to approximate native-like sucrose transport. We then compare the inferred expression scores from the scan for transmembrane helix 4 (TM4) to phylogenetic and computational thermostability analyses. As we show, caution must be applied when applying predictive methodologies to augment deep mutagenesis in the plant sciences.

A single-site saturation mutagenesis library encompassing all possible single-amino acid substitutions in AtSWEET13 was constructed using overlap extension PCR and the coding sequence for the 294 sites of the full-length protein **(Fig. 1a)**. The library was then transiently expressed in Expi293F cells under conditions that typically do not yield more than a single coding variant per cell.^11,12^ The transfected cells were incubated with 0.5 mM esculin and, after transport was quenched, stained for AtSWEET13 surface expression (via antibody staining of an extracellular N-terminal c-myc tag). Using fluorescence activated cell sorting (FACS), populations of cells expressing AtSWEET13 variants with high or low transport activity, termed AtSWEET13-High and AtSWEET13-Low sorts respectively, were collected **(Fig. S1)**. Cells that did not express AtSWEET13 variants were depleted from both sorted populations. Illumina sequencing was used to determine the frequencies of all 5880 mutants in the library using the transcripts from the naïve library and the two sorted populations. The enrichment ratio for each substitution was calculated by comparing the frequency of the transcript in the sorted population to its frequency in the naïve library.^13^ For the AtSWEET13-High sort, the enrichment ratios define the mutational landscape for enhanced cellular influx of esculin **(Fig. 1b)**. The enrichment ratios from the two independent replicates are correlated; though, agreement generally increases with variants of higher sequence frequency **(Fig. S2a-b)**. Conservation scores were calculated by averaging the log_2_ enrichment ratios at each residue and are highly correlated between the two independent replicates **(Fig. S2c-e)**.

**Fig. 1:**
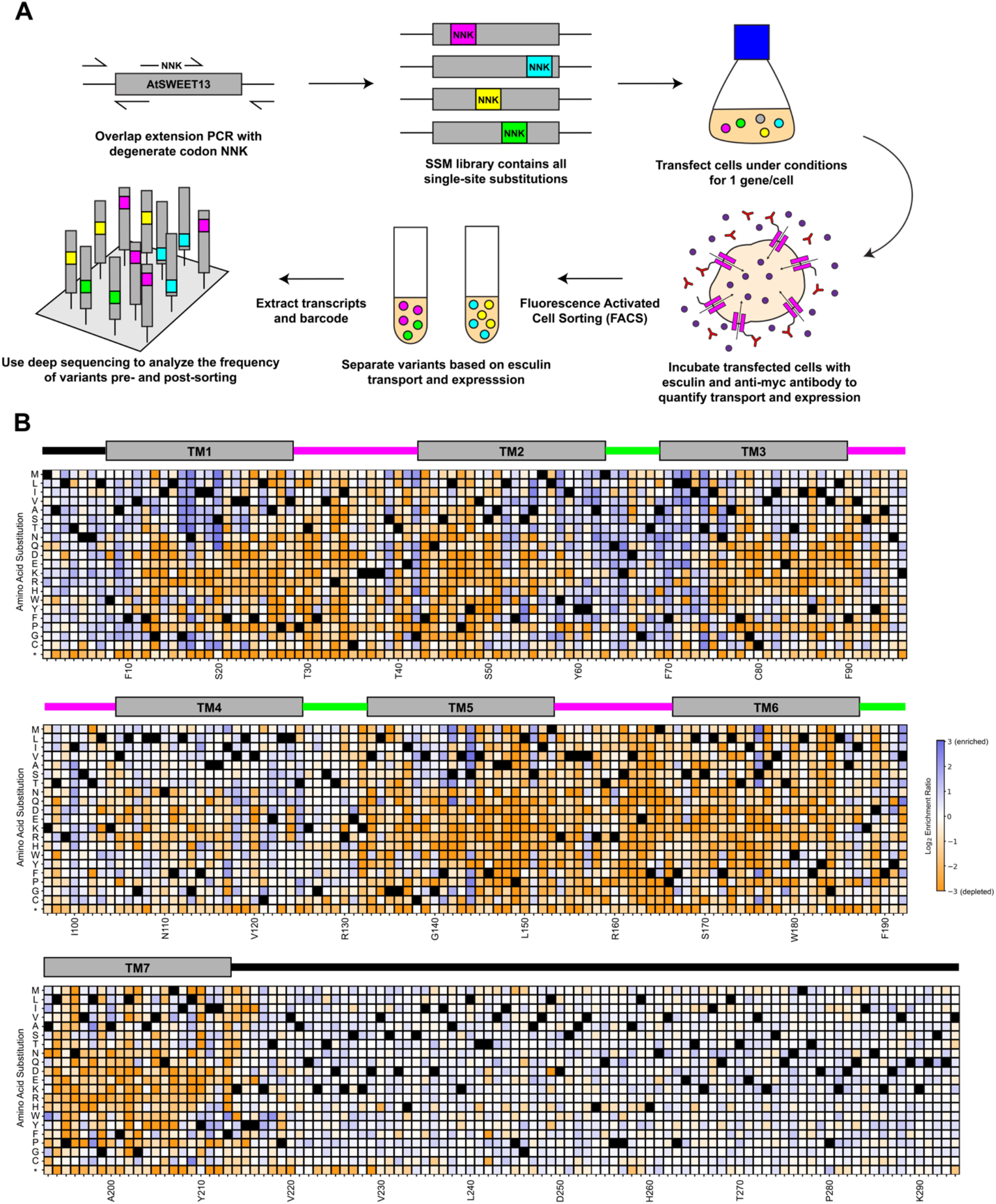
Deep mutagenesis of the AtSWEET13 sugar transporter. **a**, A schematic detailing the deep mutational scanning methodology. Using overlap extension PCR with primers encoding a degenerate codon NNK, every site in AtSWEET13 was mutagenized to all other amino acids to construct the SSM library. The library was then transfected into Expi293F cells to link cellular genotype and phenotype. Next, the cells expressing AtSWEET13 variants are incubated with esculin and anti-myc antibody and discriminated into populations based on transport and expression via FACS. After isolating the RNA transcripts from the sorted cells, both the sorted and naïve libraries were barcoded and sent to deep sequencing to determine the enrichment of variants in the different populations. **b**, Above the heat map, the secondary structure diagram of AtSWEET13 is shown, in which termini are black, transmembrane helices are grey blocks, and intra- and extracellular loops are magenta and green, respectively. The log_2_ enrichment ratios of the AtSWEET13-High sort are plotted from -3 (depleted, orange) to 0 (neutral, white) to +3 (enriched, blue). Data are averaged from two independent replicates. Wildtype amino acids are in black. The residues in the full AtSWEET13 sequence are on the horizontal axis, and the amino acid substitutions are on the vertical axis, with * as stop codons.

Membrane transporters follow an alternating access mechanism in which transition between two major conformations, the outward-facing (OF) and inward-facing (IF) states, controls substrate translocation.^14^ Intrafacial, or cytosolic, residues control gating of the former, while extrafacial, or extracellular, residues control gating of the latter. Although AtSWEET13 has only been crystallized in the IF state,^15^ both molecular dynamics (MD) simulations and site-directed mutagenesis studies of AtSWEET13 provide strong evidence for an alternating access mechanism.^16,17^ On the basis of the MD simulations of AtSWEET13, we hypothesize that shifting the conformational equilibrium of AtSWEET13 for favoring the OF state over the IF state leads to enhanced cellular influx of sugars. Mapping the conservation scores from the AtSWEET13-High sort onto the structure of AtSWEET13 bound to sucrose in an IF-like state from MD simulations reveals that the overall structure is conserved for esculin transport **(Fig. S3a)**.^18^ Enriching mutations for esculin influx are clustered at the extrafacial side around the N-terminus and the loop between TM 2 and 3, whereas the intrafacial side remains largely intolerant to substitutions especially around the loop between TMs 5 and 6 **(Fig. S3b and Fig. 2a-b)**. The cytoplasmic tail located at the C-terminus of AtSWEET13 possesses very weakly enriching mutations **(Fig. 1b and Fig. S3b)**, suggesting that the tail plays little role in enhancing esculin transport. Although C-terminal phosphorylation for other sucrose-transporting SWEETs is important for regulation of sucrose transport and drought stress responses,^19^ the specific plant protein machinery required to maintain these forms of regulation are not present in the mammalian cells utilized by this study.

**Fig. 2:**
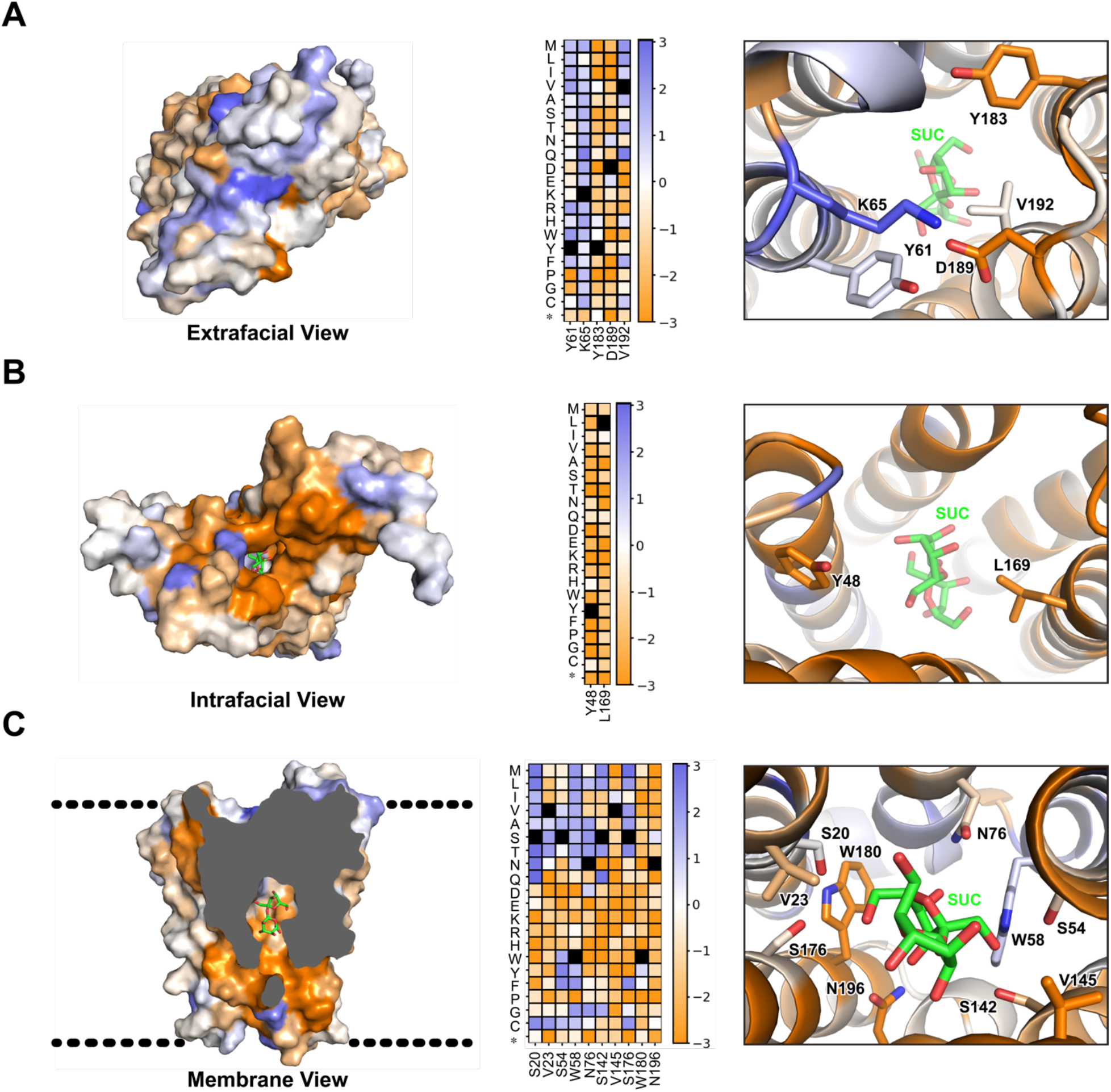
The extrafacial side of AtSWEET13 is more tolerant of mutations for enhanced esculin influx than the intrafacial side and substrate channel. Conservation scores from the AtSWEET13-High sort are mapped onto the IF-like state from MD simulations (shown as a surface) from three perspectives: extrafacial (**a**), intrafacial (**b**), and membrane (**c**) with a cross-section to show the channel. Regions where mutations are enriched or depleted for esculin transport are shown in blue or orange, respectively. Heat maps display the log_2_ enrichment ratios from the AtSWEET13-High sort, plotted from -3 (orange, depleted) to +3 (dark blue, enriched), for three sites: extracellular gating (**a**), intracellular gating (**b**), and substrate binding site (**c**). Residue positions are on the horizontal axis, while substitutions are on the vertical axis. Asterisks (*) denote stop codons. Wildtype amino acids are in black. Magnified views of the three sites are displayed on the right, and residues follow the same color scheme as the structures on the left.

The extracellular gate is comprised of AtSWEET13-Y61, K65, Y183, D189, and V192, with hydrogen bonds between the side chains of AtSWEET13-Y61, K65, and D189 stabilizing the IF state as shown in the crystal structure and the conformation derived from the molecular dynamics simulation of AtSWEET13 **(Fig. 2a)**.^15,18^ The high mutational tolerance at K65 implies it is not functionally critical for the extracellular gate and is a “hotspot” for shifting the conformational equilibrium of AtSWEET13 possibly by destabilizing the IF state. In contrast, the intracellular gate formed by Y48 and L169 is very mutationally intolerant since these residues are critical for OF state stabilization and cellular influx of sucrose via AtSWEET13 **(Fig. 2b)**. Although the interaction between these two residues is not immediately apparent in the crystal structure and the simulation state **(Fig. 2b)**, alanine substitutions at Y48 and L169 have shown to reduce and abolish sucrose transport, respectively, which correlates with their depletion in the DMS.^15^

As expected, the transport channel is highly conserved to maintain interactions for proper substrate binding and translocation **(Fig. 2c)**. In particular, the residues of the substrate binding site, aside from W58, are conserved for substrate transport, in line with their previously established functional importance.^5,15^ Notably, substitutions to charged amino acids are poorly tolerated across all residues in the substrate binding site. Replacing V23, S54, V145, and S176 with the corresponding residues from monosaccharide transporters (Leu, Asn, Met, and Asn substitutions, respectively) are deleterious, suggesting that substrate transport may be sensitive to molecular determinants for monosaccharide-selectivity. Additionally, in the substrate binding site, there appears to be an asymmetry in conservation across residues in the two triple helix bundles, despite the inverse symmetry of the residue locations. Residues in TMs 5, 6, and 7 (S142, V145, S176, N196, and W180) are overall more conserved for transport than the ones in TMs 1, 2, and 3 (S20, V23, S54, W58, and N76), indicating that TMs 5, 6, and 7 may play a more important role in recognition and transport of substrate than TMs 1, 2, and 3. MD simulations of both glucose and sucrose transport by AtSWEET13 confirm that TMs 5, 6, and 7 are critical to intracellular sugar recognition.^18^ Aside from results specific to gating and direct substrate recognition, dimensionality reduction techniques also depict generalizable interpretations of the DMS data throughout the AtSWEET13 protein sequence and structure **(Fig. S4)**. Overall, our DMS results suggest esculin can meaningfully capture native-like disaccharide transport function by AtSWEET13.

We then validated the sucrose transport capabilities of AtSWEET13 variants enriched for transporting esculin through sucrose uptake assays using a FRET biosensor **(Fig. 3)**. Five novel AtSWEET13 variants for enhanced sucrose transport were identified. AtSWEET13 and S144 is a membrane-facing residues. Therefore, the S144I variant may promote AtSWEET13 expression and enhance transport by improving membrane insertion via reduced hydrophilic character. AtSWEET13-A2 and G42 are located at the extrafacial and intrafacial gates, respectively, suggesting that A2N and G42H variants may affect gating and/or substrate recognition prior to translocation. From mutations in the binding pocket shown to directly bind sucrose in the 5XPD crystal structure, V145I was able to enhance esculin and sucrose transport. Additionally, S176M also enhanced sucrose transport. These mutations show that the characterized SWEET disaccharide-preferring “VSVS” motif possesses greater plasticity than previously thought.^15^ These assays reveal that enriched esculin transport can indeed translate to improved sucrose transport. We attempted to utilize these findings by designing higher-order mutations, but only found the combination of A2N/G42H/S144I to have greater sucrose transport activity than wildtype. Nonetheless, using a fluorescent analogue to enable cell sorting and selection also poses limitations. The S20Q variant is highly enriched in the positive esculin selection yet displays lower than wildtype sucrose transport, as previously reported.^5^ Aromatic substitutions at S54Y and S176F indicate that enhanced esculin transport may arise from aromatic stacking interactions with esculin’s coumarin ring. This exact interaction is absent from sucrose transport. Deviations in mutational landscapes can be attributed to differences in substrate perception; however, measurements of mutational fitness would not be possible in our DMS methodology without a fluorescent signal.

**Fig. 3:**
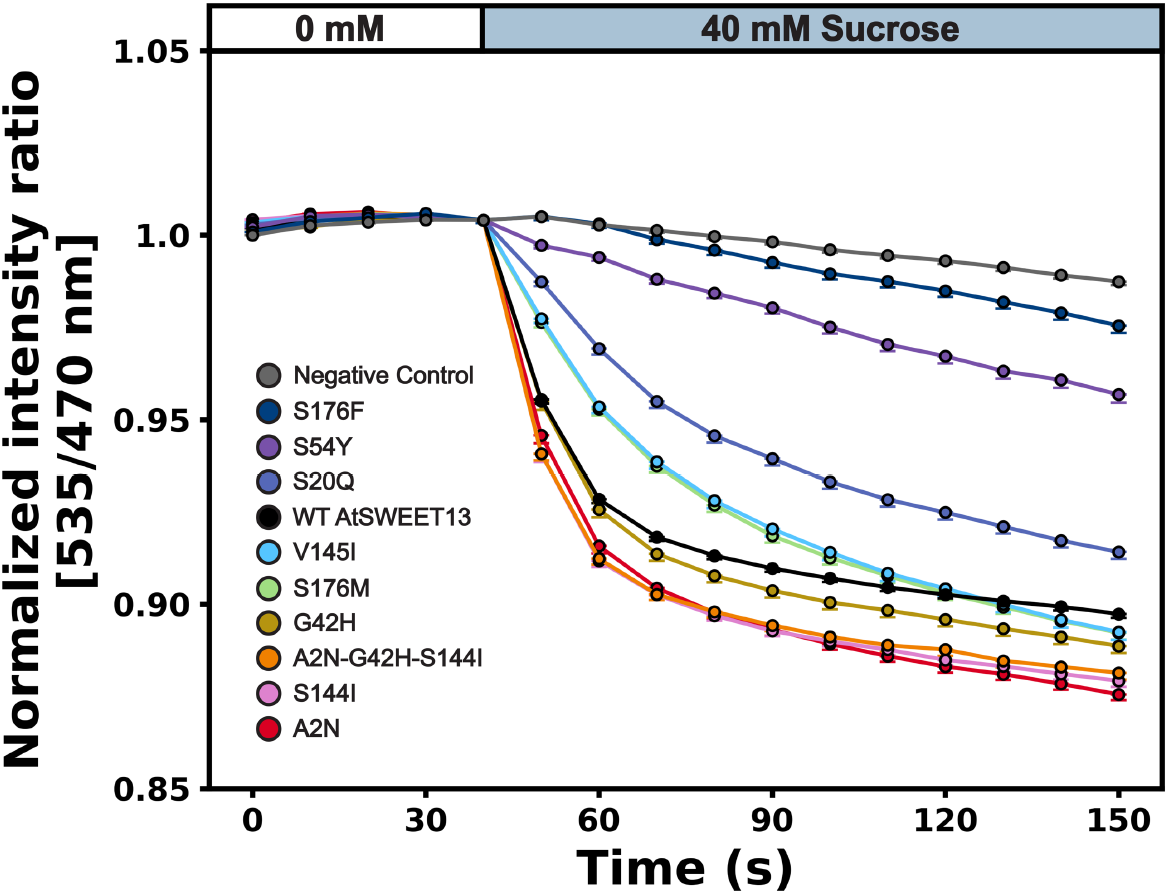
Validation of AtSWEET13 variants enriched for esculin transport using sucrose uptake. Expi293F cells co-expressing wildtype or variants of myc-tagged AtSWEET13 and sucrose FRET biosensor, FlipSuc-90u delta1, were incubated in Hank’s buffered saline solution, pH 7.4 (HBSS). At time point 40s, 40 mM sucrose in HBSS was added, and a negative ratio change was observed. Final solution concentration is equivalent to 20 mM sucrose. For the negative control, no significant change in the ratio was observed since cells expressed only the sucrose biosensor. Data are mean ± SEM, N = 3 biological replicates.

We have shown that DMS results could help identify the key mutations for enhanced transport function but they could also be used to explain evolutionary constraints on protein structure-function.^7^ Here, we explore the relationship between TM4 sequence composition throughout SWEET transporter land plant phylogeny. TM4 is the structural hallmark separating plant SWEETs from prokaryotic SemiSWEETs, comprising the seven transmembrane helical toplogy.^20,21^ A homotrimeric OsSWEET2b crystal structure exhibits TM4 as an oligomerization interface between monomeric SWEET protomers.^21,22^ MD simulations of the AtSWEET13 that resolved the full sugar transport cycle have also shown TM4 undergoes substantial fluctuations during commitment to alternate access.^18^ Overall, TM4 is integral to SWEET membrane insertion, structural integrity, and function. Considering how the four phylogenetically divided SWEET clades demonstrate different substrate preferences,^23^ perhaps clade-specific TM4 composition in light of our DMS data can reveal evolutionary divergence across clades.

A phylogenetic tree capturing ∼5000 land plant SWEET sequences was generated to provide clade-specific annotation **(Fig 4a)**. Following annotation, the sequences were aligned to AtSWEET13 per clade to identify a sequence frame corresponding to TM4. Clade-specific TM4 sequence motifs were then derived **(see Computational Methods)**.

**Fig. 4:**
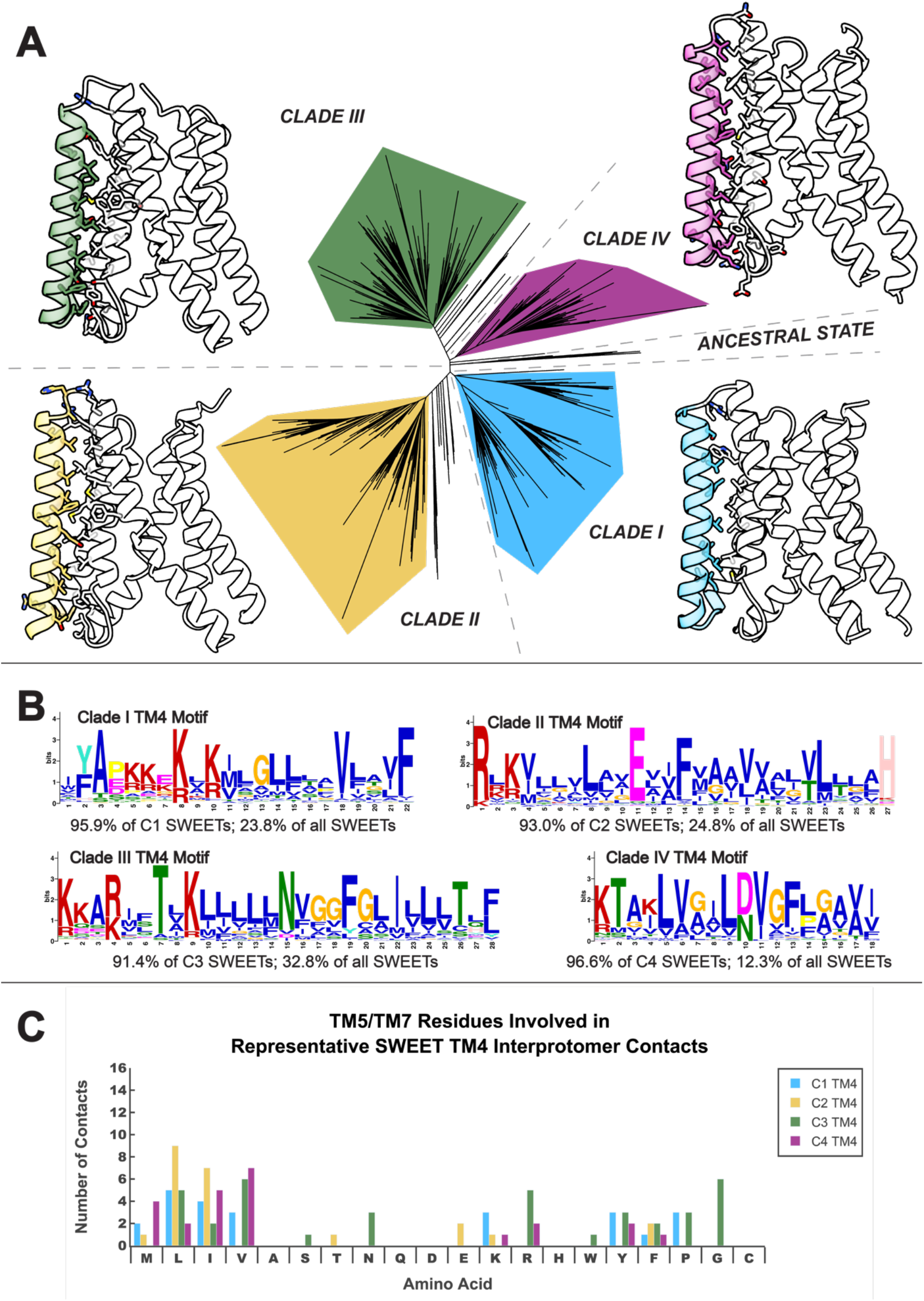
Clade-specific SWEET phylogenetic mapping reveals interhelical interactions for oligomerization. **a**, A phylogenetic tree for land plant SWEETs divided across Clades I, II, III, and IV. The structures for representative protein sequences for each clade were predicted using state-of-the-art structure prediction software. Representative sequences were chosen based on the probability of their TM4 motif provided STREME enrichment probabilities. Representative structures are modeled based off UniProt accessions A0A218VUP4 (Clade I; *Punica* granatum), A0A0A9QMY2 (Clade II; *Arundo donax*), B4FBY7 (Clade III; *Zea mays*), and A0A7J7D0X8 (*Tripterygium wilfordii*). **b**, Clade-specific TM4 sequence STREME motif logos and their prevalence within each SWEET clade. TM4 sequence windows were predicted based off multiple sequence alignment to the AtSWEET13-TM4 sequence and are highlighted on structures shown in (**a**). **c**, Interhelical contacts with non-TM4 residues occurring within a 5 Å radius along the TM4-TM5/TM7 interprotomer interface. Clade-specific results are reported based on hypothetical trimer assemblies predicted for the structures shown in (**a**).

State-of-the-art structural modeling of SWEETs from different clades suggests satisfactory putative TM4 window determination from multiple sequence alignment **(Fig. 4a)**.^24–29^ Each clade-specific motif residue favors small and aliphatic amino acids, followed by big, charged polar or aromatic amino acids **(Fig. 4b)**. These large aromatic/charged amino acids prefer outward flanking motif positions, with small/aliphatic amino acids interspersed. Such interaction types are also seen in hypothetical trimeric assemblies involving these predicted structures **(Fig. 4c)**. The same van der Waals, C/H-pi, and hydrogen bonded interactions describe intraprotomer packing within the OsSWEET2b trimer structure, while interactions with aromatic residues dominate the interprotomer TM4-TM6/7 interfaces.^22^

Universally equipped with similar physicochemistry for tight packing interactions and the same topology, one would imagine TM4 clade-specific motif substitution to be relatively tolerated. Substitutions for all possible combinatorial clade-specific TM4 motifs onto AtSWEET13 were evaluated by referencing the expression scores for the equivalent point mutation **(see Computational Methods)**. The experimental expression scores were derived by adding the log_2_ enrichment ratios from the AtSWEET13-High and -Low sorts. While this approach cannot truly reflect the complexities of epistasis, it may approximately suggest where and what AtSWEET13-TM4 substitutions are permissible.

Surprisingly, our analyses revealed TM4 in AtSWEET13 to be on average mutationally intolerant to any clade-specific substitution **(Fig. S5)**. Because the motif windows per clade are smaller than the number of residues spanning AtSWEET13-TM4, our combinatorial substitution approach permits larger probabilities for AtSWEET13 wildtype residues to be maintained or sampled along the flanking portions of TM4.

However, when TM4 positions assumed to be fixed as wildtype throughout all possible motif substitutions are excluded, the majority case is that site-specific substitution leads to AtSWEET13 single-point mutations with depleted expression. Given that AtSWEET13 belongs to Clade III, it is unsurprising that Clade III motif swapping presents the most instances where a substitution samples an expression-enhancing or wildtype mutation. Meanwhile Clade IV (fructose-transporting SWEETs), presents an overwhelming number of expression-depleting mutations in the transmembrane region. Monosaccharide-transporting Clades I and II SWEET-TM4 motifs present greater instances for more probable expression-enhancing mutations than Clade IV but are still dominated by single-point substitutions shown to deplete AtSWEET13 expression from our mutational scan. These results suggest clade-specific neofunctionalization manifests as mutational intolerance in TM4 composition. TM4 positional and compositional preferences may reinforce evolutionary constraints via tissue-specific SWEET expression and localization of specifically transported sugars.^30^

It cannot always be assumed that the results from a DMS of one protein may readily translate to another.^7^ Ideally, a computational framework could accurately predict the effects of mutation without exhaustive DMS experiments per protein of interest. For the plant sciences, this would be especially advantageous when studying homologs across different crop species. A variety of *in silico* variant prediction tools exist for evaluating mutation effect on protein thermostability and function **(see Computational Methods)**.^31–36^ Each software describes the protein using some combination of physicochemical descriptors, structural signatures, and physics-based energy terms, either with or without machine learning. Unlike for soluble proteins, the extent of amino acid hydrophobicity versus solvent accessibility does not directly correlate with functional conservation for membrane proteins. Membrane protein thermostability datasets are also less available, making force field or model optimization more prone to overfitting errors than what is already seen in tools for soluble proteins. These software are therefore known to perform better for small, soluble, globular proteins.^37^ The software used in this manuscript were selected because of their popularity or dedicated use for membrane protein analysis.

By evaluating TM4, which does not directly participate in substrate recognition, we provide a rigorous test case for determining how trustworthy *in silico* thermostability prediction software are for an example plant protein. In addition to traditional tools, we also develop our own machine learning models using state-of-the-art protein language model embeddings trained on membrane protein thermostability values **(see Computational Methods)**.^38–41^ To make software output qualitatively comparable to experiments, all results were rescaled, normalized, and analyzed as relatively (de)stabilizing across AtSWEET13-TM4 mutations.

Overall, RosettaMP tends to perform the best for relative AtSWEET13-TM4 predictions, although the performance for each software varies by the type of substitution **(Fig. S6 and S7)**. This is likely because RosettaMP implements a scoring function designed specifically for membrane proteins, which evaluates the impact of substitution associated with neighborhood side chain repacking within 8 Å of the mutation site. Accounting for changes in space nearby a mutation site has been suggested to be critical for improving predictive software accuracy.^37,42^ Reflecting the bias towards alanine scanning in training datasets, all methods incorrectly predict destabilizing mutations when going from smaller to big/aromatic residues. While RosettaMP maintains higher accuracy, some mutation results cannot be physically computed due to steric clash.^43^ RosettaMP maintains superiority for most relative aliphatic and charged substitution predictions. mCSM appears to perform the best for predicting relative uncharged polar substitution effects. mCSM, PoPMuSiC, and RosettaMP perform almost equally for predicting relatively big substitution effects. STRUM performs best for predicting relatively small substitution effects. Each of these data were *relatively* compared for results specific to AtSWEET13-TM4. We recommend using data transformation protocols so that trends towards a specific protein can be understood despite inherent software inaccuracies.

It is commonly understood that most *in silico* variant effect prediction tools reproduce generally correct trends while sacrificing absolute accuracy. These errors tend to propagate from models which overfit data-poor and biased datasets with descriptors that do not entirely account for the changes in adjacent amino acid space incurred from mutation.^37,42^ Implementing state-of-the-art language model embeddings to produce our own transfer machine learning model for AtSWEET13-TM4 prediction could not overcome these limitations, highlighting accurate prediction dependence on training data quality **(Fig. S8-S10)**. Per our phylogenetic analyses, blind reliance on *in silico* tools would have led to the interpretation that TM4 is more mutationally intolerant than our experiments show. For example, these software failed to capture TM4 accommodation for aromatic or charged polar substitutions, which would likely be used to promote oligomerization.

For plant biologists looking to use *in silico* tools to guide experiments, we recommend *qualitatively* relying on a *relative* consensus of results from appropriate software trained on data for proteins which are very similar to the target. Greater qualitative success can be achieved by accounting for how mutation changes nearby protein structure, which is generally captured in computationally expensive physics-based tools. Physics-based software often rely on available protein structures, which can be confidently predicted today **(see Computational Methods)**.^24–29^ Machine learning tools for variant effect prediction relying on evolutionary sequence profiles have also been developed, but successful implementation depends on curating sequence profiles with higher extents of homology,^44^ which is challenging for some plant proteins (like AtSWEET13 in this study). Recently developed pretrained language model embeddings can be calculated without deriving sequence profiles,^38,39^ creating opportunity for researchers to predict variant effects within and across plant proteomes. Transfer or zero-shot learning software require pre-provided DMS results for comparison, making them largely inaccessible for predicting single-point mutations within the plant sciences. These embeddings should accurately represent protein sequences and hopefully reproduce DMS datasets, provided trustworthy reference datasets for machine learning. Most current datasets are overwhelmingly bacterial or mammalian in origin, making epistasis in plants discernable a nearly impossible task. However, provided a DMS dataset, machine learning developments can theoretically enable the prediction of higher-order mutants for dramatic enhancement of desired phenotypes.^44^

Agreement with literature-reported mutations and validation of novel variants offers credence that our AtSWEET13 scan could be used for explaining variant effects on AtSWEET13 evolution and function. Marginal success was achieved for predicting relative variant effects for TM4 mutation, but we recommend how plant biologists can use *in silico* methods for soluble proteins. Application of zero-shot learning showed that current software predictions cannot match our experimental results, despite performing within the lower range of reported ESM-1v capabilities **(Fig. S11)**. However, language model embeddings can offer a chance to learn from plant protein scans for downstream applications (e.g., higher-order mutation development for engineering crops and commercial plants). Single or higher-order substitutions derived from our AtSWEET13 DMS have the potential to generate variants with enhanced expression profiles and sugar transport functions in crops.

## METHODS

### Deep Mutagenesis

The pCEP4-myc-AtSWEET13 construct was generated by cloning *Arabidopsis thaliana* SWEET13 (Genbank NP_199893.1) into the NheI-XhoI sites of pCEP4 (Invitrogen) with a N-terminal HA leader (MKTIIALSYIFCLVFA), c-myc tag, and linker (GSPGGA). A single-site saturation mutagenesis library of the full AtSWEET13 sequence was constructed using pCEP4-myc-AtSWEET13 as the template. Every position of AtSWEET13 was mutagenized using overlap extension PCR and introducing degenerate codons NNK at each site.^45^ Expi293F cells (ThermoFisher Scientific) were cultured in Expi293 Expression Medium (ThermoFisher Scientific) at 125 rpm, 8% CO_2_, and 37°C. Expi293F cells were transfected with 1 ng of library coding plasmid diluted in 1.5 µg of pCEP4-ΔCMV carrier plasmid per mL of culture at 2 × 10^6^ / mL using Expifectamine (ThermoFisher Scientific). These conditions typically yield no more than a single coding variant per cell^11,12^, linking cell genotype to phenotype. The medium was replaced 2 h pos-transfection with fresh medium, and at 24 h pos-transfection, the cells were harvested for fluorescent activated cell sorting (FACS). The cells were first washed with room-temperature PBS supplemented with 0.2% bovine serum albumin (PBS-BSA) and then incubated for 5 minutes at room-temperature with 0.5 mM esculin (Millipore Sigma, E8250). Transport of esculin was quenched with ice-cold PBS-BSA. Cells were incubated for 20 minutes on ice with anti-myc Alexa 647 (clone 9B11, 1/250 dilution, Cell Signaling Technology) in PBS-BSA. After washing twice with PBS-BSA, the cells were sorted on an Invitrogen Bigfoot Spectral Cell Sorter at Roy J. Carver Biotechnology Center. To remove dead cells, debris, and doublets, the main cell population was gated by forward/side scattering. The 15-20% of cells with the highest and the 25% with the lowest esculin fluorescence from the myc-positive (Alexa 647) population were collected in tubes **(Fig. S1)**, prepared by coating overnight with fetal bovine serum and adding Expi293 Expression Medium prior to sorting. Total RNA was extracted from the collected cells using a GeneJET RNA purification kit (ThermoFisher Scientific), and complementary DNA (cDNA) was reverse transcribed with Accuscript (Agilent) primed with a gene-specific oligonucleotide. The full mutagenized sequence of AtSWEET13 was PCR amplified as five fragments. A second round of PCR was performed on the fragments to add adapters for unique barcoding, annealing to Illumina sequencing primers, and binding to the flow cell. The PCR products were sequenced on an Illumina NovaSeq 600 using 2 × 250 nucleotide paired end protocol, and the resulting data were analyzed using Enrich.^13^ The log_2_ enrichment ratios for each AtSWEET13 variant was calculated using the frequencies of variants in the transcripts of sorted populations and their frequencies in the naïve plasmid library. The ratios were then normalized by subtracting the log_2_ enrichment ratio for the wildtype sequence. Conservation scores at each site were calculated by averaging the log_2_ enrichment ratios for all nonsynonymous mutations at the same site.

### Sucrose Uptake Assay

The gene encoding the sucrose biosensor for fluorescence resonance energy transfer (FRET), FlipSuc-90u delta1, was subcloned from pRSET FlipSuc-90u delta1 into the BamHI-HindIII sites of pcDNA3.1(+). pRET FlipSuc-90u delta1 was a gift from Wolf Frommer (Addgene plasmid #14475; http://n2t.net/addgene:14475; RRID: Addgene_14475). To measure sucrose uptake of AtSWEET13 wildtype and variants, Expi293F cells were co-transfected with 500 ng of pcDNA3.1(+)-FlipSuc-90u delta1 and 500ng of pCEP4-myc-AtSWEET13 (wildtype or mutants) per mL of culture at a density of 2*10^6^ cells/mL using Expifectamine (ThermoFisher Scientific). For the negative control, Expi293F cells were transfected with only 500 ng of pcDNA3.1(+)-FlipSuc-90u delta1 per mL of culture at a density of 2*10^6^ cells/mL using Expifectamine (ThermoFisher Scientific). At 5 h post-transfection, the cells were centrifuged to remove media containing transfection reagents and resuspended in fresh Expi293 Expression Medium (ThermoFisher Scientific). Resuspended cells were then seeded onto a 96-well plate coated with Poly-D-lysine (Gibco) and containing fresh culture media. At 16 h post-transfection, the cells were imaged at intervals of 10s for 210s using a Zeiss AxioObserver Z1/7 with a Hamamatsu camera ORCA-Flash4.0 V3Digital CMOS (exposure time of 50 ms), X-Cite XYLIS LED Light (CFP excitation was at 430 nm with 20nm bandwidth, YFP excitation was at 470/40), a 40x dry objective lens and the Hamamatsu W-VIEW GEMINI splitting optics (emission from each excitation was split and captured to two channels, CFP at 480/40 and YFP at 535/30).^46^ Culture media in the sample plates were replaced with 50 ul Hank’s buffered saline solution, pH 7.4 (HBSS, prepared as done previously),^47^ followed by adding 50 ul 40 mM sucrose in HBSS at time point 100s. The final solution concentration is equivalent to 20 mM sucrose. The 210s ratio change (delta ratio) of CFP excitation-YFP emission intensity divided by CFP excitation-CFP emission intensity for cells was recorded using ZEN 3.5. Three biological replicates were performed for AtSWEET13 wildtype and variants and for the negative control. For each biological replicate, six technical replicates were performed by collecting the delta ratio of 10 cells for each technical replicate. Data points on curves were delta ratio from 60s to 210s, and error bars were standard error mean.^5^ Mutants and wildtype curves were shifted to better align with the negative control curve before the addition of sucrose at 40s.

Computational methods are discussed in detail in the Supplementary Information.

## DATA AVAILABILITY

Illumina sequencing data are deposited in NCBI’s Gene Expression Omnibus (GEO) under series accession number ######. Codes and files used in this manuscript have been made available via our github repository: https://github.com/ShuklaGroup/AtSWEET13_DMS

## ACKNOWLEDGEMENTS

D.S. acknowledges support from the National Institute of Health (Award No. R35GM142745). Staff at the UIUC Roy J. Carver Biotechnology Center assisted with FACS and Illumina sequencing. We thank Christine A. Devlin for assisting with computational tools for preparation of heat maps. We thank Lars Kruse for helpful advice concerning phylogenetic tree bootstrapping. We thank Nathalie Vercruysse and Professor Marianne Rooman at the Université libre de Bruxelles for authorizing our private account request for use of Dezyme’s PoPMuSiC software. We thank Nicole Chiang for helpful conversations concerning data normalization, rescaling, and dimensionality reduction.

## Author Contributions

D.S. acquired funding for this project. D.S. conceived and supervised the project. K.K.N., E.P. and D.S. performed deep mutational scanning experiments and analyses. A.T.W. performed phylogenetic and structural bioinformatics analyses and conducted dimensionality reduction techniques on DMS data. L.X. optimized and performed FRET sucrose sensor assays and analyzed assay results. X.M. generated and optimized machine learning models, as well as executed zero-shot predictions. L.-Q.C. and C.Z. provided laboratory materials and helped optimize and troubleshoot AtSWEET13 expression and FRET assays. E.P. helped optimize and troubleshoot AtSWEET13 deep mutational scanning experiments, as well as provide feedback on deep mutational scanning results and phylogenetic analyses. K.K.N., A.T.W., L.X. and X.M. wrote the manuscript with inputs from D.S., E.P., and L.-Q.C. Authors K.K.N., A.T.W. and L.X. contributed equally. All authors approve of the final version of this manuscript.

## ETHICS DECLARATION

There are no competing interests.

## SUPPLEMENTAL INFORMATION

This letter contains Supplemental Information. Supplemental Information contains additional experimental and computational results, as well as a thorough description of all computational methods.

