## Supplementary Information for "Deep mutational scanning reveals sequence to function constraints for SWEET family transporters"

### Table of Contents


**Computational Methods** S-3

**Figure S1** Gating strategy for discriminating AtSWEET13 variants based on expression and esculin transport S-14

**Figure S2** Agreement between two independent replicates of the AtSWEET13 deep mutational scan S-15

**Figure S3** The mutations at the N-terminus and the loop between TMs 2 and 3 are enriched for esculin influx, while those at the loop between TMs 5 and 6 is highly depleted S-16

**Figure S4** Data-driven dimensionality reduction reveals AtSWEET13 residue clusters with generalizable perturbation profiles S-17

**Figure S5** Combinatorial clade-specific substitution reveals AtSWEET13-TM4 to be mutationally intolerant. S-18

**Figure S6** Measuring how well can *in silico* thermostability and variant effect prediction software reproduce AtSWEET13-TM4 DMS data – Part 1 S-19

**Figure S7** Measuring how well can *in silico* thermostability and variant effect prediction software reproduce AtSWEET13-TM4 DMS data – Part 2 S-20

**Figure S8** Performance of different regression models on the MPTherm training dataset (n=917) S-21

**Table S1** Hyperparameter search for SVM, RF, and MLP models training on ProSE embeddings for the MPTherm dataset S-22

**Figure S9** Evaluating performance of ProSE transfer learning models against AtSWEET13-TM4 data – Part 1 S-23

**Figure S10** Evaluating performance of ProSE transfer learning models against AtSWEET13-TM4 data – Part 2 S-24

**Figure S11** ESM-1v zero shot learning is unable to accurately reproduce AtSWEET13 DMS data S-25

### COMPUTATIONAL METHODS

#### *Curating a dataset of land plant SWEET transporter sequences*

The curation of SWEET land plant sequences began with downloading the dbSWEET repository and verifying the sequences in the UniProt database.[1,2] The organism list provided in the dbSWEET repository was re-queried in UniProt and expanded by using the search keywords "SWEET bidirectional transporter". UniProt sequences lacking experimental annotation were retained only if their sole PFAM identifier was PF03083 (MtN3_slv family; sugar efflux transporters for intercellular exchange). A keyword search for "SWEET bidirectional transporter" was repeated in the Joint Genome Institute's Phytozome database,[3] where land plant sequences were retained. The combined list of UniProt and Phytozome sequences were provided to the HHblits webserver offered within the Max Planck Bioinformatics Toolkit to identify additional hits for sequence retrieval not previously captured during UniProt and Phytozome searches.[4,5] In total, this manually curated land plant SWEET dataset contains ~5000 sequences. A multiple sequence alignment (MSA) was made from the final list of land plant SWEET sequences with the MAFFT Version 7 Large Alignment Server using the L-large-INS-1 option with all other default settings present.[6] ClipKIT trimming software was used to clean highly divergent sites in the MAFFT-derived MSA as a final preparative step before phylogenetic tree construction.[7]

The trimmed MAFFT MSA of all ~5000 SWEET sequences served to generate an initial maximum likelihood tree using RaxML v8.2.[8] Trial runs using the PROTOGAMMAIAUTO flag were performed to determine the highest scoring amino acid substitution model for the SWEET MSA, ultimately yielding the JTT amino acid substitution model as the best. A maximum likelihood tree was constructed with the following command:

*/path/standard-RAxML-master/raxmlHPC-PTHREADS-AVX -T 64 -m PROTGAMMAIJTTX -n raxml_tree.phy -p 12345 -s mafft_clipkit_msa.fasta*

Due to computational cost, bootstrapping of the RAxML-derived tree proceeded using the ultrafast bootstrapping option in IQ-TREE.[9–11] Multicore bootstrapping was performed using the following command:

*/path/iqtree -s mafft_clipkit_msa.fasta -st AA -m JTT+I+G4+FO+ASC -bb 1000 -wbt -wbtl -alrt 1000 -abayes -t raxml_tree.newick -wt -nt AUTO -keep-ident -pre iqtree_ufbs_bootN*

Where *iqtree_ufbs_bootN* reflected the multicore bootstrap replicate. Across 20 multicored replicates, a total of 29000 bootstraps were performed on the RAxML guide maximum-likelihood tree. From these bootstraps, a consenus tree was prepared using the following command:

```
/path/iqtree -con -t alltrees -nt AUTO
```

Lastly, bootstrap support values were calculated using the following command:

```
/path/iqtree -sup alltrees.contree.trees -t alltrees -nt AUTO -keep-ident
```

In total, for all SWEET sequences, 95.56% have a bootstrap support value of 100; 1.01% have support values between 90 and 100; 0.14% have support values between 80 and 90; 3.15% have support values between 70 and 80; and 0.14% have support values between 60 and 70. Identifying clades as SWEET Clades I-IV was determined based on the localization of *Arabidopsis thaliana* SWEET sequences throughout the tree. As a safety check, the positions of other experimentally characterized SWEETs were in agreement with the SWEET Clade assignment.[12]

#### *Motif enrichment analysis throughout phylogenetic tree-labeled clades*

After successful differentiation of SWEET clades through the consensus phylogenetic tree, each of the curated SWEET sequences were separated by clade. Each clade was then aligned to AtSWEET13 using similar methods as described above. Based on sequence alignment to AtSWEET13 and visualization of AtSWEET13 structures,[13,14] a sequence region was estimated to equate to transmembrane helix 4 (TM4) across the different clades. Each of the clade-specific fasta sequences were then trimmed where each fasta entry would only include the supposed TM4 region. Any sequence entries where gaps existed along this TM4 stretch were pruned for motif enrichment analysis using STREME.[15] STREME analyses within the MEME Biosuite were performed using the following advanced settings.[16] For clade-specific motif analyses, the query input involved a single clade; meanwhile, a fasta file containing all other clades was used as a control. For a motif search across all SWEETs, a single fasta was queried containing all characterized clades with a control option using randomized sequences. The motif width was up to 28 amino acids, the length of the TM4 helix in the AtSWEET13 crystal structure. The p-value threshold was set to 0.05. The Markov order was set to '0' where motifs were arranged by left ends. Resulting sequence logos and results were downloaded.

#### *Combinatorial motif substitution along AtSWEET13-TM4*

The purpose of this analysis is to approximate the likelihood of whether AtSWEET13-TM4 can accommodate clade-specific single-point mutations that are expected to be seen across the TM4 sequence compositions expressed in land plant SWEETs. Due to the complexities of epistasis when accumulating higher order mutations, it is unlikely that this analysis can truly evaluate the ability for clade-specific TM4 motifs to be exactly swapped. However, this substitution analysis can show, given an individual point mutation that is sampled by SWEETs within a specific clade, whether our single-point mutagenesis data reveals that exact substitution to enrich or deplete AtSWEET13 expression. Alternatively, an involved single-point mutation could also sample an AtSWEET13-TM4 wildtype residue. All combinatorial possible sequence motifs for each SWEET clade were enumerated, given that the amino acid types present in the motif revealed STREME probabilities of greater than or equal to 10%. Because the potential regions for substitution of clade-specific motifs onto AtSWEET13-TM4 could be variable, a motif scanning analysis was performed. Sliding windows were generated where each of the possible motifs could be then evaluated as to whether each of the given point mutations resulted in enhanced or depleted expression or sampled a wild-type amino acid residue. The "hypothetical" placement of SWEET-TM4 motifs was determined based on alignment of polar residues at the beginning of each to the polar residues in AtSWEET13-TM4. Namely, agreement with Lys96, Lys97, Arg99, and Lys104. For demonstrative purposes, the positioning of each clade-specific TM4 motif onto AtSWEET13-TM4 is illustrated below. Any instance where a digit is enumerated instead of a letter indicates where an amino acid sampled within a clade-specific TM4 motif was introduced.

For Clade I:

**WT** **A N K K T R I S T L K V L G L L N F L G F A A I V L V C E L L T K**
**C1-1** 1 1 1 1 1 1 1 1 1 1 1 1 1 1 1 1 1 1 1 1 1 1 A I V L V C E L L T K
**C1-2** A 1 1 1 1 1 1 1 1 1 1 1 1 1 1 1 1 1 1 1 1 1 1 I V L V C E L L T K
**C1-2** A N 1 1 1 1 1 1 1 1 1 1 1 1 1 1 1 1 1 1 1 1 1 1 V L V C E L L T K
**C1-4** A N K 1 1 1 1 1 1 1 1 1 1 1 1 1 1 1 1 1 1 1 1 1 1 L V C E L L T K

For Clade II:

**WT** **A N K K T R I S T L K V L G L L N F L G F A A I V L V C E L L T K**
**C2-1** 2 2 2 2 2 2 2 2 2 2 2 2 2 2 2 2 2 2 2 2 2 2 2 2 2 2 V C E L L T K
**C2-2** A 2 2 2 2 2 2 2 2 2 2 2 2 2 2 2 2 2 2 2 2 2 2 2 2 2 2 C E L L T K
**C2-2** A N 2 2 2 2 2 2 2 2 2 2 2 2 2 2 2 2 2 2 2 2 2 2 2 2 2 2 E L L T K
**C2-4** A N K 2 2 2 2 2 2 2 2 2 2 2 2 2 2 2 2 2 2 2 2 2 2 2 2 2 2 L L T K

For Clade III:

**WT** **A N K K T R I S T L K V L G L L N F L G F A A I V L V C E L L T K**
**C3-1** A N 3 3 3 3 3 3 3 3 3 3 3 3 3 3 3 3 3 3 3 3 3 3 3 3 3 3 3 L L T K

For Clade IV:

**WT** **A N K K T R I S T L K V L G L L N F L G F A A I V L V C E L L T K**
**C4-1** A N 4 4 4 4 4 4 4 4 4 4 4 4 4 4 4 4 4 4 F A A I V L V C E L L T K
**C4-2** A N K 4 4 4 4 4 4 4 4 4 4 4 4 4 4 4 4 4 4 A A I V L V C E L L T K

Results were plotted using a dot plot variant of a stacked bar graph using the Plotly python library.[17]

#### *Representative SWEET protein structure predictions*

Structural placement of bioinformatically predicted TM4 regions was verified using state-of-the-art structure prediction software.[18–22] Representative SWEET sequences were selected based on the STREME probability calculated for the incorporation of their respective TM4 sequence motifs. Because resolved structures of trimeric SWEET transporters depict each protomer as homoprotomers (i.e., each SWEET participating in the complex was resolved in the same conformation),[23] hypothetical SWEET trimers were arranged by structurally aligning each of the clade-specific representative predicted structures with each protomer chain obtained from the OsSWEET2b timer crystal structure (PDB ID: 5CTH). Interhelical contacts were then evaluated via visual analysis by comparing TM4 residue – non-TM4 residue contacts occurring within a 5Å radius. Results were plotted as a bar graph using Plotly.[17]

#### *In silico mutagenesis – Input PDB file preparation*

Based off preexisting simulation data which resolved the conformational transitions and transport cycle for AtSWEET13 sucrose transport, metastable states were identified with respect to molecular recognition of sucrose. Based on sucrose positioning throughout transport,[13] six states were identified as potential inputs for *in silico* thermostability prediction software. Approximately 100 representative structures were extracted from each metastable state given the preexisting molecular dynamics simulation dataset and reformatted as PDB structures using CPPTRAJ. To improve computational throughput on *in silico* prediction measures, pairwise root-mean-square deviation (RMSD) calculations were performed using the backbone atom selections @CA,C,O,N in Chimera command line to find the most representative structure (i.e., lowest average pairwise RMSD) from the energetic minima.[24] The average pairwise RMSD between structures per metastable state was approximately 2 Å or less.

#### *Rosetta Membrane Protein ΔΔG calculations*

For Rosetta calculations, the representative PDB files were cleaned (i.e., renumbering all atoms and residues from ‘1’, replacing all hydrogens and nonstandard amino acids with names interpretable by Rosetta software). Using the C implementation of the Rosetta 2022-332 precompiled binary software release for Linux, a membrane spanfile was generated.[25] The purpose of the spanfile is to assert what parts of a protein structure are transmembrane spanning. The dimensions of the spanfile were refined given *a priori* knowledge of AtSWEET13 packing into a model membrane bilayer using the CHARMM-GUI Membrane Builder.[26,27] PyRosetta calculations for implicit membrane insertion, structure minimization, and *in silico* mutagenesis were then performed using membrane protein (MP) methods.[25,28,29] A pip wheel for installing a Python 3.6 compatible version of PyRosetta 2022 was acquired from Rosetta Commons. The PyRosetta session was initialized using a few extra options. pH mode was enabled where the pH was set to 7.4 to match experimental conditions. The implicit membrane settings were set such that no preexisting pore was assumed to be present. The membrane thickness value (equivalent to half the bilayer thickness) was set to 10.5 Å based on CHARMM-GUI outputs. Out of the options available in PyRosetta, the lipid composition was set to DOPC (18:1/18:1) as to best approximate the average lipid tail unsaturation seen in a model plant plasma membrane bilayer.[30] Multithreading was enabled. Lastly, the franklin2019 scoring function was used based on its specificity for membrane protein-related calculations.[31] In agreement with preestablished protocols for soluble proteins in the Rosetta ddg_monomer application, the AtSWEET13 PDB structures were minimized following a low resolution protocol.[32] Harmonic restraints were applied to only the Cα atoms.[32] A MPFastRelax minimization was then performed to help relax the AtSWEET13 PDB structures while embedded in an implicit DOPC bilayer. After minimization, the wildtype residues along TM4 (A94 – K126) were mutated using the `mutate_residue` command from the PyRosetta toolbox. In accordance with the “low-resolution” protocol used in ddg_monomer calculations for soluble proteins, mutations were accompanied by rotamer repacking of nearby residue sidechains within an 8 Å radius of the designated mutation site with no further minimization.[32] All mutations were scored using the franklin2019 scoring function.[31] In agreement with the ddg_monomer protocol, each residue was mutated to every possible amino acid 50 times, where the average of the top three most stable values was recorded. The MP_ΔΔG of mutation was then reported as the difference between the $\Delta G^{mutation}$ minus the $\Delta G^{WT}$. $\Delta G^{WT}$ is equivalent to the score obtained by using the `mutate_residue` function to reintroduce the wildtype amino acid at its own position.

*Webserver-based thermostability calculations*
The Biosig Lab mCSM-membrane webserver was provided with each PDB structure and a mutation list corresponding to every possible amino acid substitution along the AtSWEET13-TM4 residues.[33] For dezyme's PoPMuSiC webserver, a private account was created to permit uploading of private PDB files obtained from molecular dynamics simulation.[34–36] Systematic mutation thermostability calculations were performed using PoPMuSiC. For both mCSM-membrane and PoPMuSiC calculations, PDB structures were supplied that were either (1) directly obtained from molecular dynamics simulation or (2) minimized using the PyRosetta minimization protocol for membrane proteins. STRUM webserver systematic results were determined solely through input of the AtSWEET13 UniProt sequence.[37] All webserver results were downloaded manually.

#### *Using ProSE language model embeddings to predict variant effects*

The MPTherm database was downloaded and formatted for training machine learning models to predict changes in melting temperature ($\Delta T_m$) as a result of mutations to AtSWEET13-TM4.[38] Each mutant sequence was recorded in fasta format and then transformed using the ProSE language model-based protein sequence embeddings.[39] ProSE embeddings were performed using the pretrained prose_mt model with an average pooling. For each input sequence, ProSE generates a 6165-dimensional representation for each amino acid. The average of all amino acid representations was used as a final embedding for each sequence (6165-dimensional vector per sequence). The embeddings of 917 membrane protein fasta entries from the MPTherm database were fed into downstream training models. In this work, we trained Support Vector Machine Regressor (SVM), Random Forest Regressor (RF) and Multilayer Perceptron (MLP) to predict the $\Delta T_m$ (°C) of membrane proteins. For SVM and RF, we used Principal Component Analysis (PCA) to reduce the dimensionality of the raw embeddings from 6165 to 35 to explain ~95% of data variance. Hyperparameters were optimized through a grid search and cross validation (Table S1). R2, RMSE, Spearman correlation and Pearson correlation were used to evaluate the performance of each model. After optimizing hyperparameters, the entire MPTherm dataset was used to refit all these three models with optimized hyperparameter selection. Then the trained models were transferred to predict the $\Delta T_m$ (°C) of the AtSWEET13-TM4 mutants. Scikit-learn was used to implement used to implement SVM, RF, and MLP.[40] R2, RMSE, as well as Spearman and Pearson correlation coefficients were calculated using the SciPy python package.[41]

#### *In silico mutagenesis data processing*

*In silico* mutagenesis data were parsed into csv files using bash scripting. The results of *in silico* calculations are largely dependent on the quality of the input. Given how conformational states representing different stages of AtSWEET13 sucrose transport were available based on molecular simulation results, we proceeded with two different types of data analysis. Firstly, following the textbook biochemical principle of energetic coupling, we added the mutation scores across each of the states representing different transport states into a single sum. The idea here is that the effect of a mutation on protein stability would be incorporated into all stages of transport, which are represented by each of the PDBs used for calculations. Secondly, we examined each individual PDB structure, as predicted stability scores do vary based on input structure. It is likely that future plant scientists performing these calculations would not have access to more than a single crystal structure, if any at all. Thus, comparing how results correlate between the experimental dataset and calculations from individual PDB structures is important. Results across each of the different computationally conformations were evaluated consistently. In lieu of a crystal structure, we recommend that high-quality predicted structures be for purposes other than visualization after (1) knowledge-based confirmation of predicted structure accuracy to the target protein family topology and (2) minimization using some type of short molecular dynamics simulation. Structure prediction works best when proteins with similar presumed topologies have structures already resolved. Current state-of-the-art and publicly available protein structure prediction software for retrieving predicted structures include DeepMind's AlphaFold2,[18] the Baker Lab's RoseTTaFold,[19] the Peng Lab's OmegaFold,[22] Facebook Research's ESMFold,[20,21] and the Zhang Lab's D-I-TASSER.[42] At the time of writing, these software can be easily accessed using a Google Collab notebook or the appropriate webserver :

AlphaFold2
https://colab.research.google.com/github/sokrypton/ColabFold/blob/main/AlphaFold2.ipynb

RoseTTaFold
https://colab.research.google.com/github/sokrypton/ColabFold/blob/main/RoseTTAFold.ipynb

OmegaFold
https://colab.research.google.com/github/sokrypton/ColabFold/blob/main/beta/omegafold.ipynb

ESMFold
https://colab.research.google.com/github/sokrypton/ColabFold/blob/main/ESMFold.ipynb

D-I-TASSER
https://zhanggroup.org/D-I-TASSER/

Individual amino acids can generally present among various subtypes, where the mutational permissiveness at each position reflects the functional and structural character of the amino acid under question.[43] To simplify the interpretation of deep mutagenesis and *in silico* thermostability prediction data were averaged across several amino acid groupings: Aromatic (F, Y, W); Aliphatic (A, V, I, L, M); Uncharged Polar (S, T, N, Q);

Charged Polar (R, H, K, D, E); Big (W, R, Y, H, F, N, D, K, Q, P, E, M); and Small (G, V, A, T, I, C, L, S).

For the effectiveness of *in silico* mutagenesis software on predicting stable predictions along AtSWEET13-TM4 to be evaluated, the *in silico* data needed to be compared to the experimental expression scores. However, the data presented within this paper is expressed across different units depending on the methodology. While raw output was compared to experimental expression scores, data transformation and scaling were required to offer a common scale for analysis. Predictions from mCSM-membrane, PoPMuSiC, and STRUM are described in kcal/mol. Meanwhile, PyRosetta MP_ΔΔG calculations are described in arbitrary Rosetta Energy Units (REU). Results from ProSE downstream predictions are expressed in melting temperature shifts ($\Delta T_m$). Firstly, the data were rescaled to make all data values positive. For subtype-averaged data, the minimum value, *m*, was identified within each respective method dataset (i.e., all reported values regardless of amino acid subtype). Each corresponding data point per *in silico* dataset was then transformed by adding *m*+1, such that the lowest possible value across all subtypes in each *in silico* dataset became '1' and all data points would become positive. Then, each of the positively shifted data values were transformed along a $\log_2$ scale, where the lowest datapoint would now equal zero. The experimental expression data were already prepared on a $\log_2$ ratiometric scale, so only *m* was added to each experimental data point, making the lowest possible experimental value '0'. For each dataset, a MaxAbsScaler scaling approach was performed where the values within each subtype method-specific dataset were then rescaled from '0' to '1'. Rescaling was performed by dividing each reported value by the maximum $\log_2$ per subtype method-specific dataset.

A different normalization approach was applied to ProSE machine learning model outputs because of how the units were expressed in terms of $\Delta T_m$ (°C). In general, a more positive $\Delta T_m$ score indicates that the protein has become more thermostable after mutation. So, to match the same relative scaling as was done for all *in silico* tools with results expressed in kcal/mol, the signs of all predicted $\Delta T_m$ values were first inverted. After sign inversion, the same data normalization and rescaling procedures as described above were repeated.

#### *Using zero-shot prediction with ESMFold-1v to recapitulate DMS data*

Five ESM-1v models were used to do zero-shot variant prediction for the AtSWEET13 DMS dataset.[20,44] Spearman and Pearson correlation coefficients were calculated between the ESM-1v predicted score and the experimentally reported expression score and conservation scores.

#### *Dimensionality reduction and residue cluster labeling of DMS transport data*

Unsupervised machine learning can be used to reduce complex and highly dimensional data to better extract meaningful insights in a more human-interpretable form. Dimensionality reduction techniques have previously been applied to deep mutational scanning datasets.[45] Dimensionality reduction methods work by performing a type of transformation onto input data, whereby the resulting operation results in an abstraction that allows for similar data points to be grouped more easily through "labeling". However, deriving appropriate labels during exploratory analysis for a deep mutational scanning dataset can be challenging. Each residue on AtSWEET13 has 20 possible mutations, and each mutation demonstrates a different functional response to mutation. Appropriate label assignment must balance consideration for each possible mutation as well as the ensemble mutational profile at a protein residue site. To bin the deep mutational scan transport data into appropriate categories for labeling, density-based hierarchical clustering (HDBSCAN) was implemented.[46] Density-based hierarchical clustering uses an agglomerative approach where not all data points are assigned to a cluster. In order to assign a label to every protein residue from the mutational scan, the cluster labels from HDBSCAN were interpreted and then manually assigned across all data points. HDBSCAN initially binned transport scores as follows:

[-5.0, -2.0), [-2.0, -1.0), [-1.0, -0.5), [-0.5, -0.3), [-0.3, -0.1), [-0.1, 0.0), [0.0, 0.00X), [0.00X, 0.1), [0.1, 0.3), [0.3, 0.4), [0.4, 0.5), [0.5, 0.6), [0.6, 0.9), [0.9, 1.1), and [1.1, 5.0]

To simplify downstream dimensionality reduction, we grouped these bins identified through exploratory data-driven analysis to a simplified set of following six bins:

[-5, -2), [-2, -1), [-1, -0.5), [-0.5, 0.0), [0.0, 0.5), [0.5, 1.1), and [1.1, 5]

AtSWEET13 residue sites were labeled by whether their respective conservation scores for transport activity fell within one of the six bins from our simplified set. Following label assignment, biochemical interpretations of the perturbation profiles for the residue sites belonging to each bin were made.

With our bins identified, we then applied the popular data dimensionality reduction algorithms Uniform Manifold Approximation and Projection (UMAP), as well as t-SNE, to the transport scores of our DMS dataset.[47,48] Other algorithms also exist for dimensionality reduction also exist, for those interested.[49,50] Pseudo-hyperparameter optimization was performed to yield the most aesthetic data transformation for improved visualization with our six cluster labels. We refer to this process as a "pseudo" optimization protocol because there is no exact ground truth for pursuit; hyperparameter selection is solely for visualization purposes. To improve interpretability of data presentation, the full AtSWEET13 protein sequence was obtained from UniProt (Protein Accession ID: Q9FGQ2) and graphically represented as a snake diagram with Protter.[51] The Protter output was then modified in Adobe Illustrator to correspond residue coloring with colors selected for UMAP and t-SNE scatterplot labeling.

#### COMPUTATIONAL METHODS REFERENCES


1. Gupta, A. & Sankararamakrishnan, R. dbSWEET: An integrated resource for SWEET superfamily to understand, analyze and predict the function of sugar transporters in prokaryotes and eukaryotes. *J. Mol. Biol.* **430**, 2203–2211 (2018).
2. UniProt Consortium. UniProt: The universal protein knowledgebase in 2021. *Nucleic Acids Res.* **49**, D480–D489 (2021).
3. Goodstein, D. M. *et al.* Phytozome: A comparative platform for green plant genomics. *Nucleic Acids Res.* **40**, D1178-1186 (2012).
4. Gabler, F. *et al.* Protein sequence analysis using the MPI bioinformatics toolkit. *Curr. Protoc. Bioinformatics.* **72**, e108 (2020).
5. Zimmermann, L. *et al.* A completely reimplemented MPI bioinformatics toolkit with a new HHpred server at its core. *J. Mol. Biol.* **430**, 2237–2243 (2018).
6. Katoh, K., Rozewicki, J. & Yamada, K. D. MAFFT online service: Multiple sequence alignment, interactive sequence choice and visualization. *Brief. Bioinform.* **20**, 1160–1166 (2019).
7. Steenwyk, J. L., Buida, T. J., Li, Y., Shen, X.-X. & Rokas, A. ClipKIT: A multiple sequence alignment trimming software for accurate phylogenomic inference. *PLoS Biol.* **18**, e3001007 (2020).
8. Stamatakis, A. RAxML version 8: A tool for phylogenetic analysis and post-analysis of large phylogenies. *Bioinformatics* **30**, 1312–1313 (2014).
9. Pattengale, N. D., Alipour, M., Bininda-Emonds, O. R. P., Moret, B. M. E. & Stamatakis, A. How many bootstrap replicates are necessary? in *Research in Computational Molecular Biology* (ed. Batzoglou, S.) 184–200 (Springer, 2009). doi:10.1007/978-3-642-02008-7_13.
10. Nguyen, L.-T., Schmidt, H. A., von Haeseler, A. & Minh, B. Q. IQ-TREE: A fast and effective stochastic algorithm for estimating maximum-likelihood phylogenies. *Mol. Biol. Evol.* **32**, 268–274 (2015).
11. Hoang, D. T., Chernomor, O., von Haeseler, A., Minh, B. Q. & Vinh, L. S. UFBoot2: Improving the Ultrafast Bootstrap Approximation. *Mol. Biol. Evol.* **35**, 518–522 (2018).
12. Xue, X., Wang, J., Shukla, D., Cheung, L. S. & Chen, L.-Q. When SWEETs turn tweens: Updates and perspectives. *Annu. Rev. Plant. Biol.* **73**, 379–403 (2022).
13. Weigle, A. T. & Shukla, D. AtSWEET13 transporter discriminates sugars by selective facial and positional substrate recognition. 2022.10.12.511964 Preprint at https://doi.org/10.1101/2022.10.12.511964 (2022).
14. Han, L. *et al.* Molecular mechanism of substrate recognition and transport by the AtSWEET13 sugar transporter. *Proc. Natl. Acad. Sci. U. S. A.* **114**, 10089–10094 (2017).
15. Bailey, T. L. STREME: Accurate and versatile sequence motif discovery. *Bioinformatics* **37**, 2834–2840 (2021).
16. Bailey, T. L., Johnson, J., Grant, C. E. & Noble, W. S. The MEME Suite. *Nucleic Acids Res.* **43**, W39–W49 (2015).
17. Plotly Technologies Inc. Collaborative data science. (2015).
18. Jumper, J. *et al.* Highly accurate protein structure prediction with AlphaFold. *Nature* **596**, 583–589 (2021).

37. Quan, L., Lv, Q. & Zhang, Y. STRUM: Structure-based prediction of protein stability changes upon single-point mutation. *Bioinformatics* **32**, 2936–2946 (2016).
38. Kulandaisamy, A., Sakthivel, R. & Gromiha, M. M. MPTherm: Database for membrane protein thermodynamics for understanding folding and stability. *Brief. Bioinform.* **22**, 2119–2125 (2021).
39. Bepler, T. & Berger, B. Learning the protein language: Evolution, structure, and function. *Cell Syst.* **12**, 654-669.e3 (2021).
40. Scikit-learn: Machine learning in Python - scikit-learn 1.2.1 documentation.
41. Virtanen, P. *et al.* SciPy 1.0: Fundamental algorithms for scientific computing in Python. *Nat. Methods* **17**, 261–272 (2020).
42. Zheng, W. *et al. Integrating deep neural network models with I-TASSER for accurate protein structure prediction*. https://zhanggroup.org/D-I-TASSER/help.html (2022).
43. Dunham, A. S. & Beltrao, P. Exploring amino acid functions in a deep mutational landscape. *Mol. Syst. Biol.* **17**, e10305 (2021).
44. Meier, J. *et al.* Language models enable zero-shot prediction of the effects of mutations on protein function. 2021.07.09.450648 Preprint at https://doi.org/10.1101/2021.07.09.450648 (2021).
45. Jones, E. M. *et al.* Structural and functional characterization of G protein–coupled receptors with deep mutational scanning. *eLife* **9**, e54895 (2020).
46. Campello, R. J. G. B., Moulavi, D. & Sander, J. Density-based clustering based on hierarchical density estimates. in *In: Pei, J., Tseng, V.S., Cao, L., Motoda, H., Xu, G. (eds) Advances in Knowledge Discovery and Data Mining. PAKDD 2013. Lecture Notes in Computer Science.* (Springer).
47. McInnes, L., Healy, J. & Melville, J. UMAP: Uniform manifold approximation and projection for dimension reduction. Preprint at https://doi.org/10.48550/arXiv.1802.03426 (2018).
48. van der Maaten, L. & Hinton, G. Visualizing data using t-SNE. *J. Mach. Learn. Res.* **9**, 2579 (2008).
49. Amid, E. & Warmuth, M. K. TriMAP: Large-scale dimensionality reduction using triplets. Preprint at https://doi.org/10.48550/arXiv.1910.00204 (2019).
50. Wang, Y., Huang, H., Rudin, C. & Shaposhnik, Y. Understanding how dimension reduction tools work: An empirical approach to deciphering t-SNE, UMAP, TriMAP, and PaCMAP for data visualization. Preprint at https://doi.org/10.48550/arXiv.2012.04456 (2020).
51. Omasits, U., Ahrens, C. H., Muller, S. & Wollscheid, B. Protter: Interactive protein feature visualization and integration with experimental proteomic data. *Bioinformatics* **30**, 884–886 (2014).

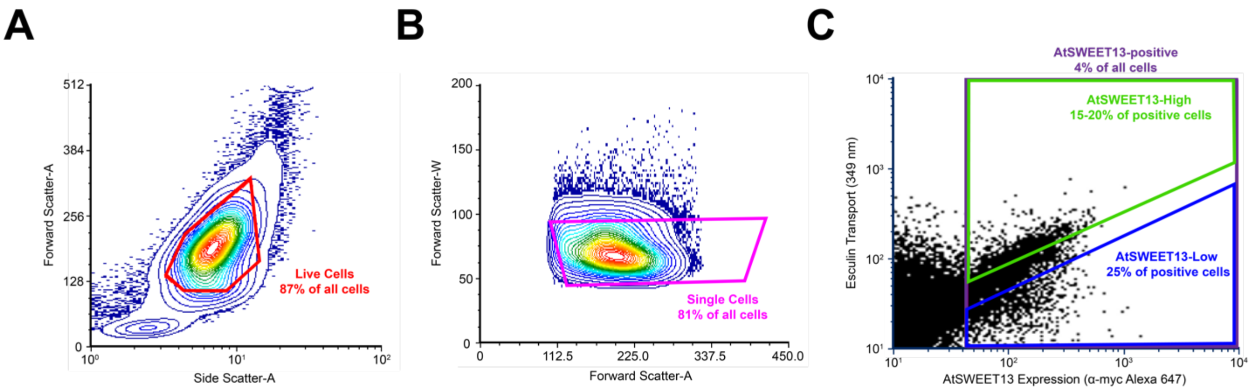


**Figure S1. Gating strategy for discriminating AtSWEET13 variants based on expression and esculin transport.** Expi293 cells transfected with a AtSWEET13 SSM library were incubated with 0.5 mM esculin for five minutes at room temperature before FACS sorting. Cells were gated for viability (**a, red**) and single cells (**b, magenta**) based on forward and side scattering to exclude dead cells, debris, and doublets from the sorted populations. The upper 15-20 and lower 25 percentiles for esculin fluorescence in AtSWEET13-expressing cells (**c, purple**) were collected in the AtSWEET13-High (**c, green**) and AtSWEET13-Low (**c, blue**) sorts.

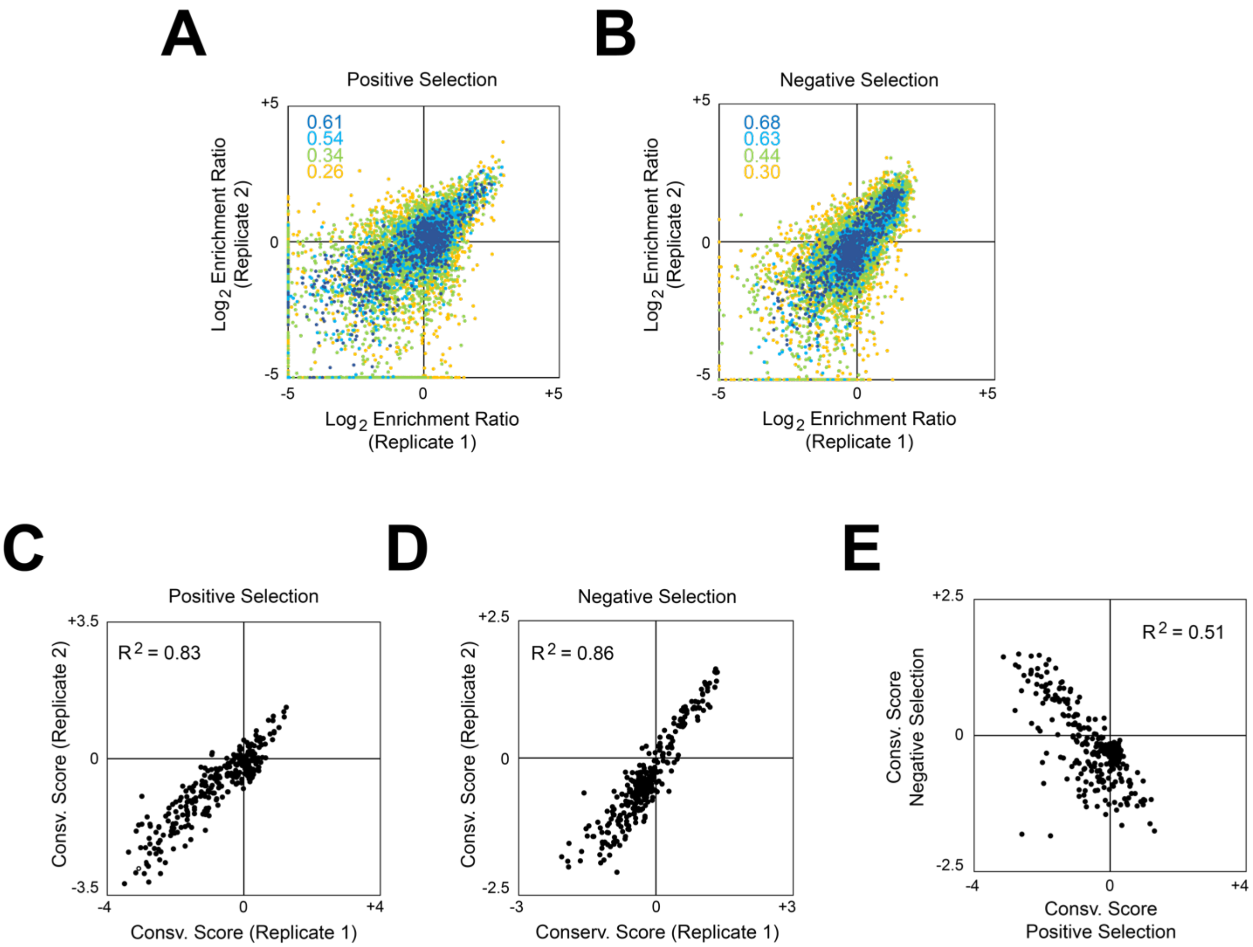


**Figure S2. Agreement between two independent replicates of the AtSWEET13 deep mutational scan. a-b,** Agreement between $\log_2$ enrichment ratios for AtSWEET13 mutations in the AtSWEET13-High (**a**) and AtSWEET13-Low (**b**) sorts, which generally increases with higher frequencies of sequence variants (blue, frequency ≥ 0.0003; cyan, 0.0003 > frequency ≥ 0.0002; green, 0.0002 > frequency ≥ 0.0001; orange, frequency > 0.0001). $R^2$ values are correspondingly colored in the top left quadrant. **c-e**, Residue conservation scores were calculated by averaging the $\log_2$ enrichment ratios for all substitutions at each position and are well correlated between two independent FACS experiments for the AtSWEET13-High (**c**) and -Low (**d**) sorts. The conservation scores between the AtSWEET13-High and -Low sorted populations (**e**) are anti-correlated. A shift towards the negative quadrant is indicative of mutations that are depleted from both sorted populations possibly due to decreased AtSWEET13 expression.

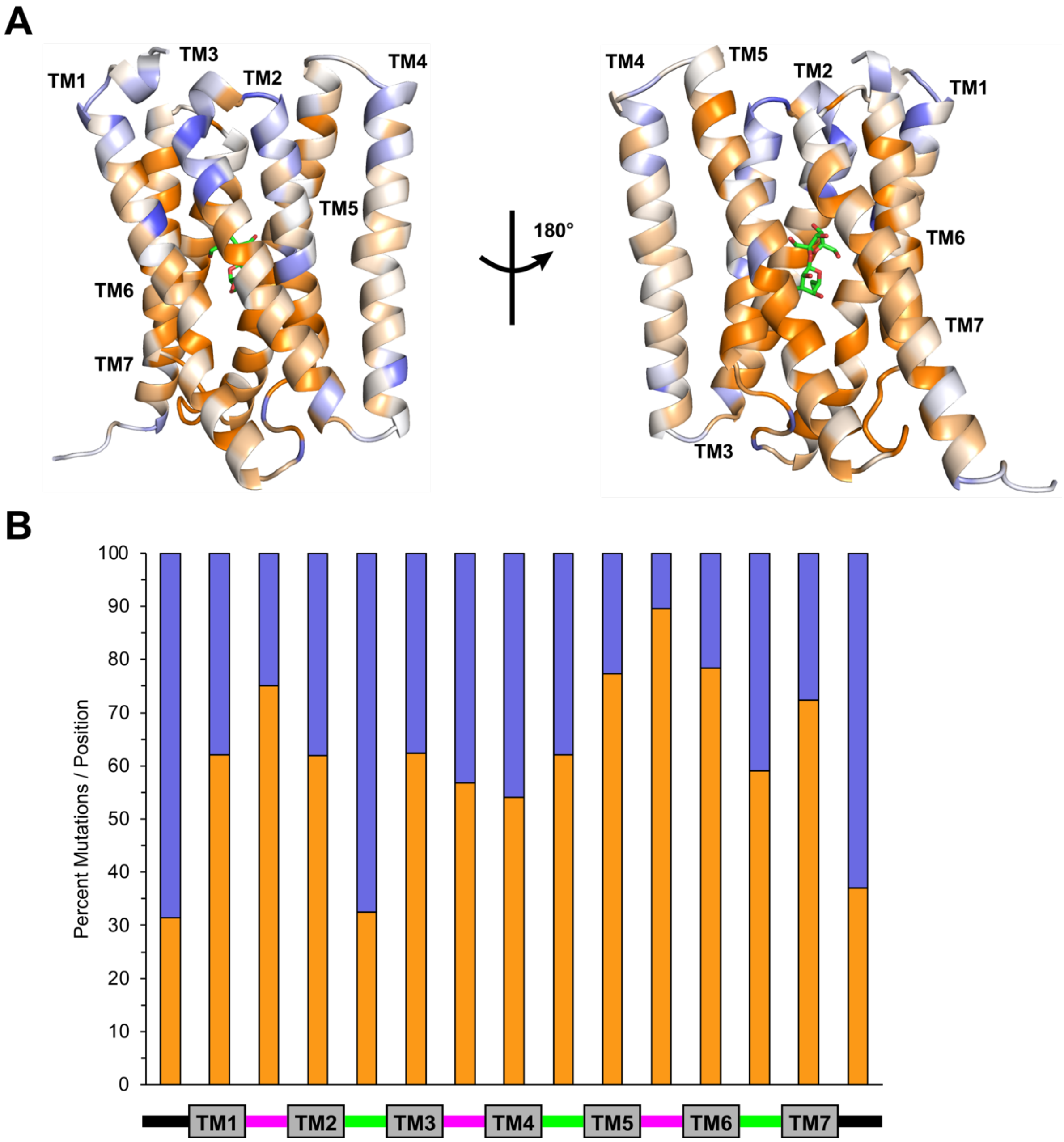


**Figure S3. The mutations at the N-terminus and the loop between TMs 2 and 3 are enriched for esculin influx, while those at the loop between TMs 5 and 6 is highly depleted. a,** Conservation scores from the AtSWEET13-High sort are mapped onto the IF-like state from MD simulations (shown as a cartoon) and rotated 180° across the z-axis. Regions where mutations are enriched or depleted for esculin transport are shown in blue or orange, respectively. **b**, The percent of enriching ($log_2$ enrichment ratio > 0, blue bars) and depleting ($log_2$ enrichment ratio < 0, orange bars) nonsynonymous mutations per total in residues in each region of AtSWEET13 are calculated and shown as a stacked bar plot. The secondary structure of AtSWEET13 is represented as a schematic below the plot with the termini (black), loops (intracellular in magenta, extracellular in green), and transmembrane helices (grey boxes).

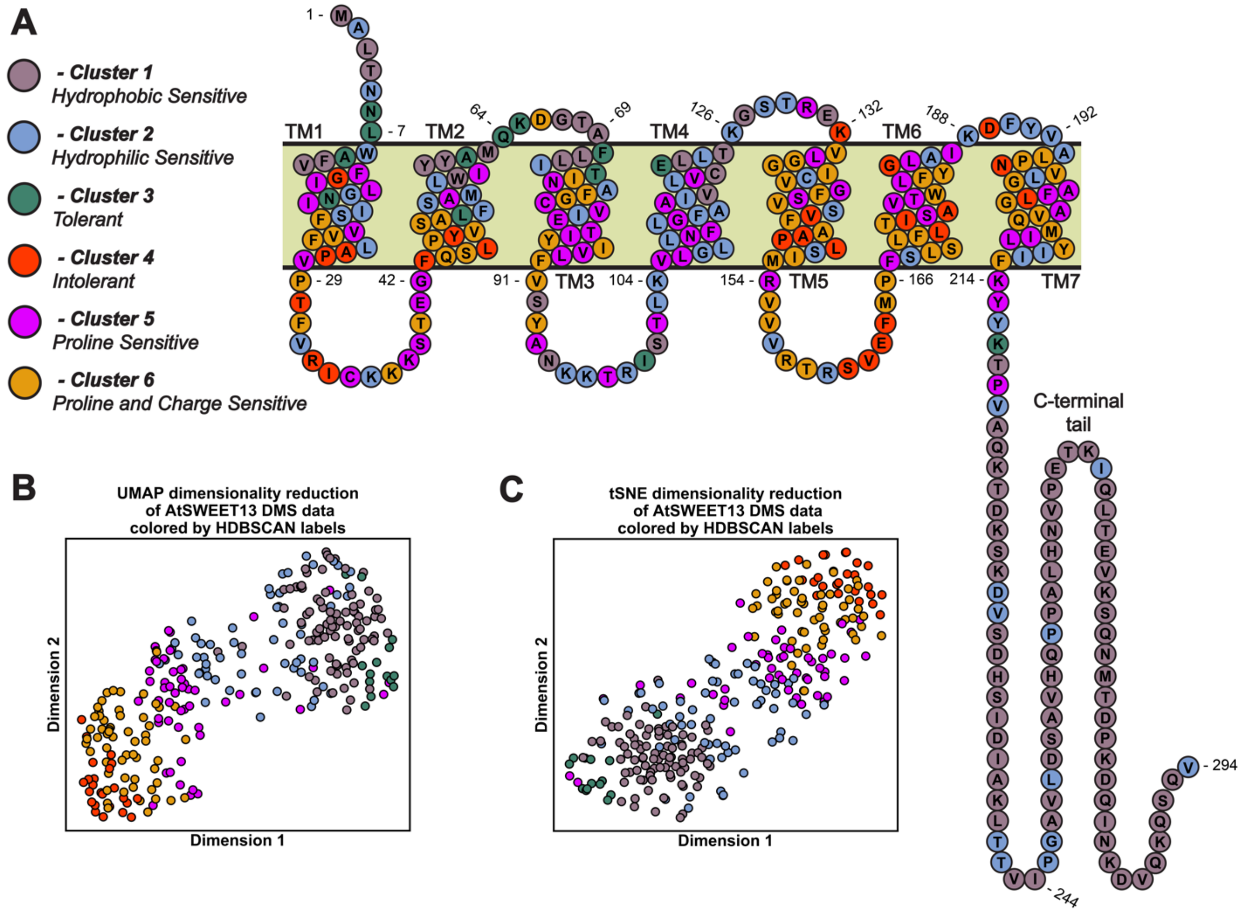


**Figure S4. Data-driven dimensionality reduction reveals AtSWEET13 residue clusters with generalizable perturbation profiles. a**, Cluster labels, their biochemical interpretation and representation onto a snake plot diagram of AtSWEET13. Cluster 1 (hydrophobic sensitive) is colored as mountbatten pink; Cluster 2 (hydrophilic sensitive) is colored as vista blue; Cluster 3 (tolerant) is colored as viridian; Cluster 4 (intolerant) is colored as coquelicot red; Cluster 5 (proline sensitive) is colored as phlox pink; and Cluster 6 (proline and charge sensitive) is colored as gamboge yellow. **b**, UMAP and **c**, t-SNE representations of AtSWEET13 transport data from the deep mutational scan.

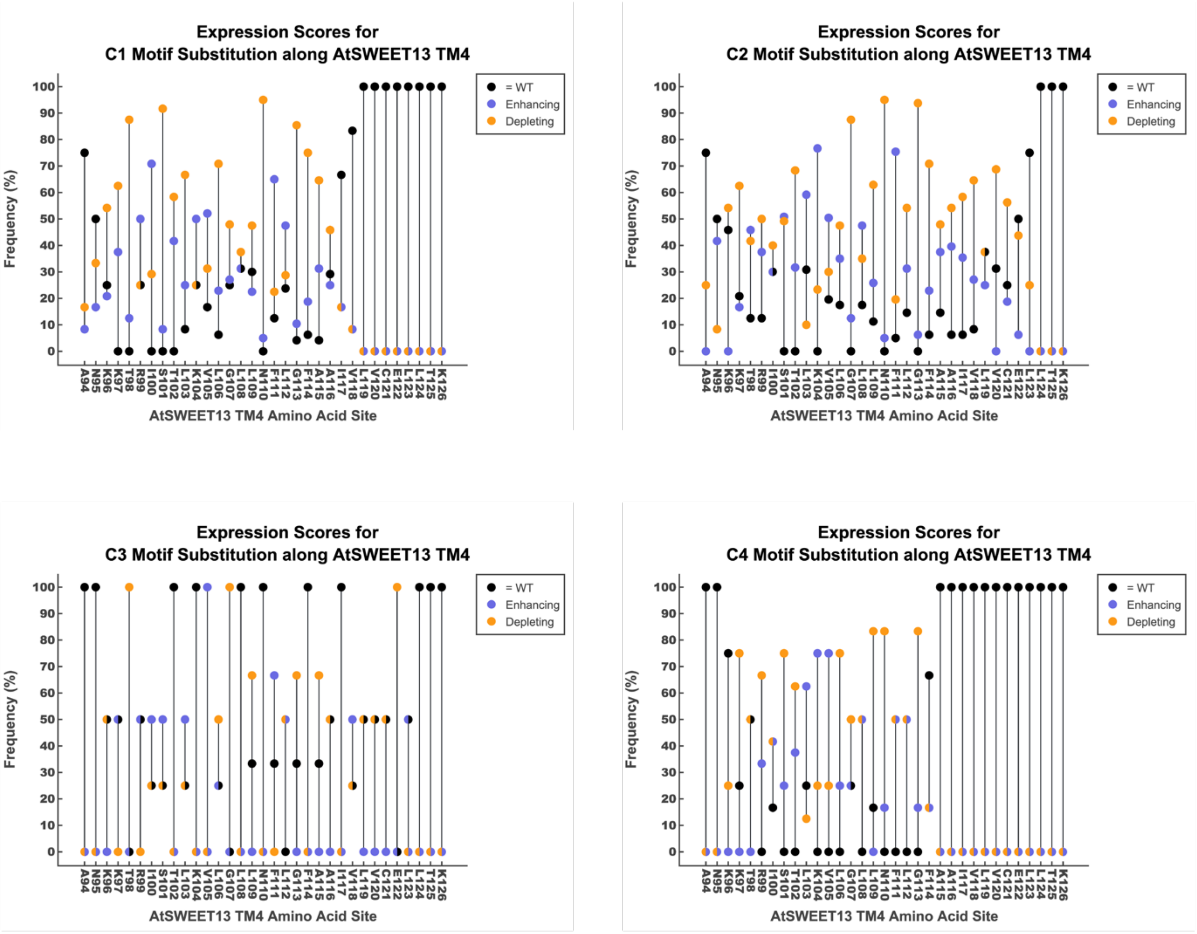


**Figure S5. Combinatorial clade-specific substitution reveals AtSWEET13-TM4 to be mutationally intolerant.** Permissible AtSWEET13-TM4 combinatorial clade-specific motif substitution per AtSWEET13-TM4 amino acid site are shown as stacked dot plots. The color of the dot positioned at the highest frequency per amino acid site indicates what is the effect of a single-point mutation enumerated in the swapped clade-specific TM4 motif. Single-point mutations are evaluated on whether the substituted motif results in enhanced (blue dots) or depleted (orange dots) expression scores based off DMS experimental results, or if the substituted motif sampled the wild-type AtSWEET13-TM4 residue (black dots). Instances where differently evaluated substitutions exhibit identical frequencies are depicted with semicircles for improved visibility. Window size for motif substitution reflects the lengths for motifs shown in Main Text Figure 4. Top left, Clade I motif substitutions (out of 477,757,440 possible motif swaps); top right, Clade II motif substitutions; bottom left (out of 2,579,890,176,000 possible motif swaps), Clade III motif substitutions (out of 764,411,904 possible motif swaps); bottom right, Clade IV motif substitutions (out of 1,327,102 possible motif swaps).

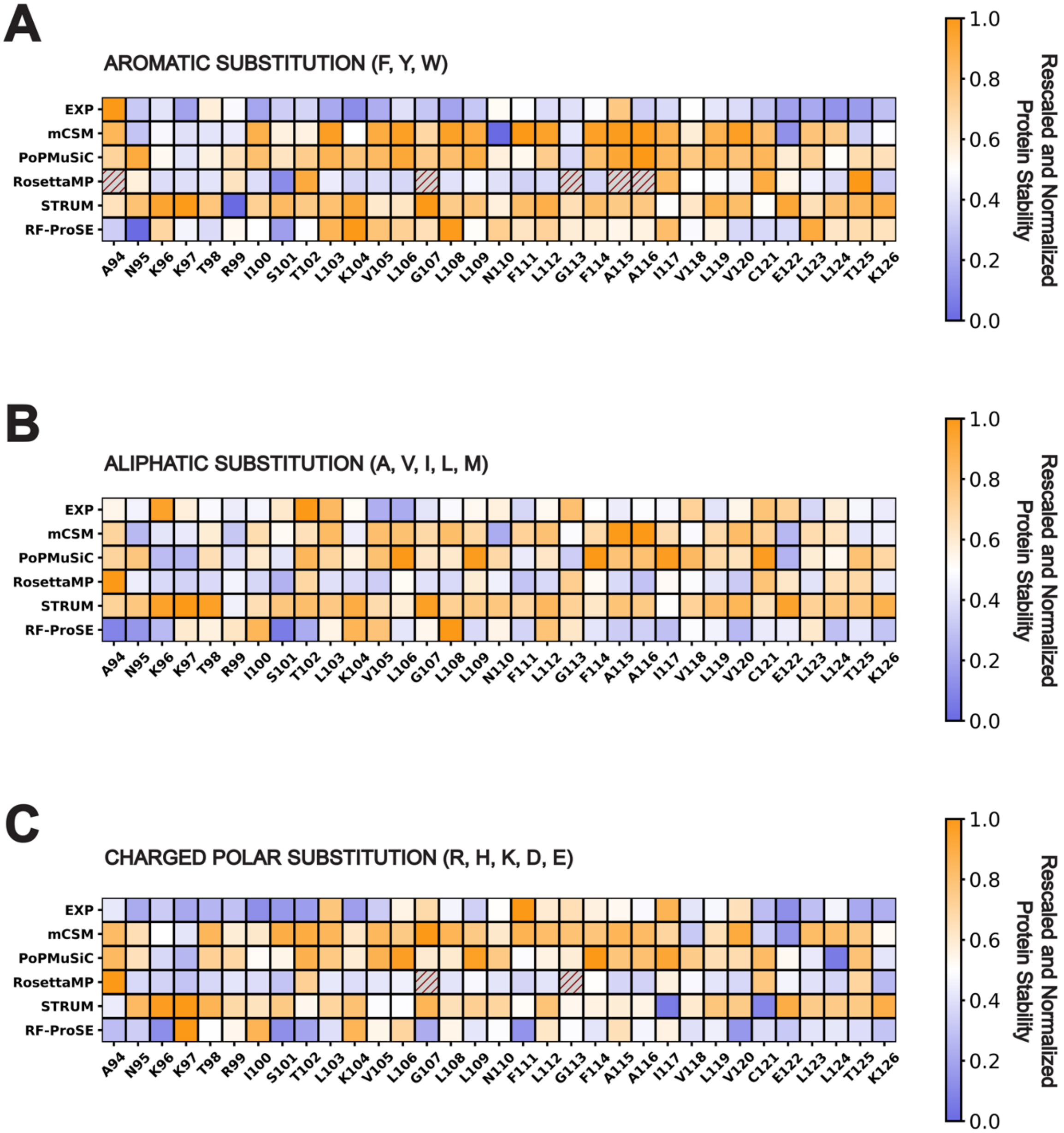


**Figure S6. Measuring how well can *in silico* thermostability and variant effect prediction software reproduce AtSWEET13-TM4 DMS data – Part 1.** Rescaled and normalized protein stability values incurred upon mutation are shown for experimental values (EXP) versus *in silico* predictions. The unit scale is expressed from 0 to 1. A value of 0 (dark blue) indicates enhanced predicted thermostability/experimentally reported expression values in response to mutation. A value of 1 (orange) indicates depleted predicted thermostability/experimentally reported expression values in response to mutation. Grey boxes with a red diagonal line pattern indicate that the *in silico* software failed to report a predicted mutation score for all of the possible amino acids enumerated in the substitution category. **a**, Aromatic amino acid substitutions (F, Y, W)**. b,** Aliphatic amino acid substitutions (A, V, I, L, M). **c,** Charged polar amino acid substitutions (R, H, K, D, E).

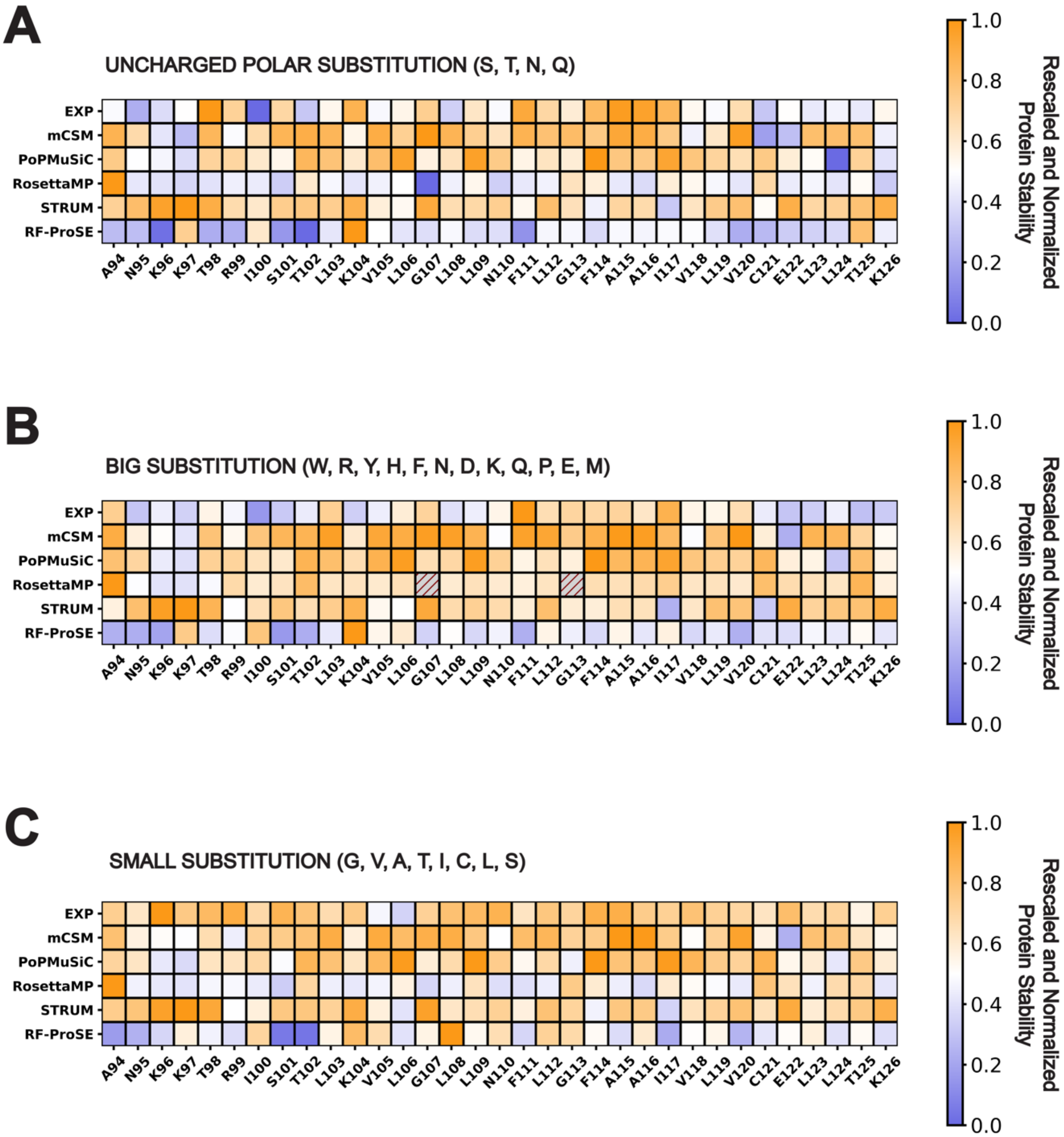


**Figure S7. Measuring how well can *in silico* thermostability and variant effect prediction software reproduce AtSWEET13-TM4 DMS data – Part 2.** Rescaled and normalized protein stability values incurred upon mutation are shown for experimental values (EXP) versus *in silico* predictions. The unit scale is expressed from 0 to 1. A value of 0 (dark blue) indicates enhanced predicted thermostability/experimentally reported expression values in response to mutation. A value of 1 (orange) indicates depleted predicted thermostability/experimentally reported expression values in response to mutation. Grey boxes with a red diagonal line pattern indicate that the *in silico* software failed to report a predicted mutation score for all of the possible amino acids enumerated in the substitution category. **a**, Uncharged polar amino acid substitutions (S, T, N, Q)**. b,** Big amino acid substitutions (W, R, Y, H, F, N, D, K, Q, P, E, M). **c,** Small amino acid substitutions (G, V, A, T, **I**, C, L, S).

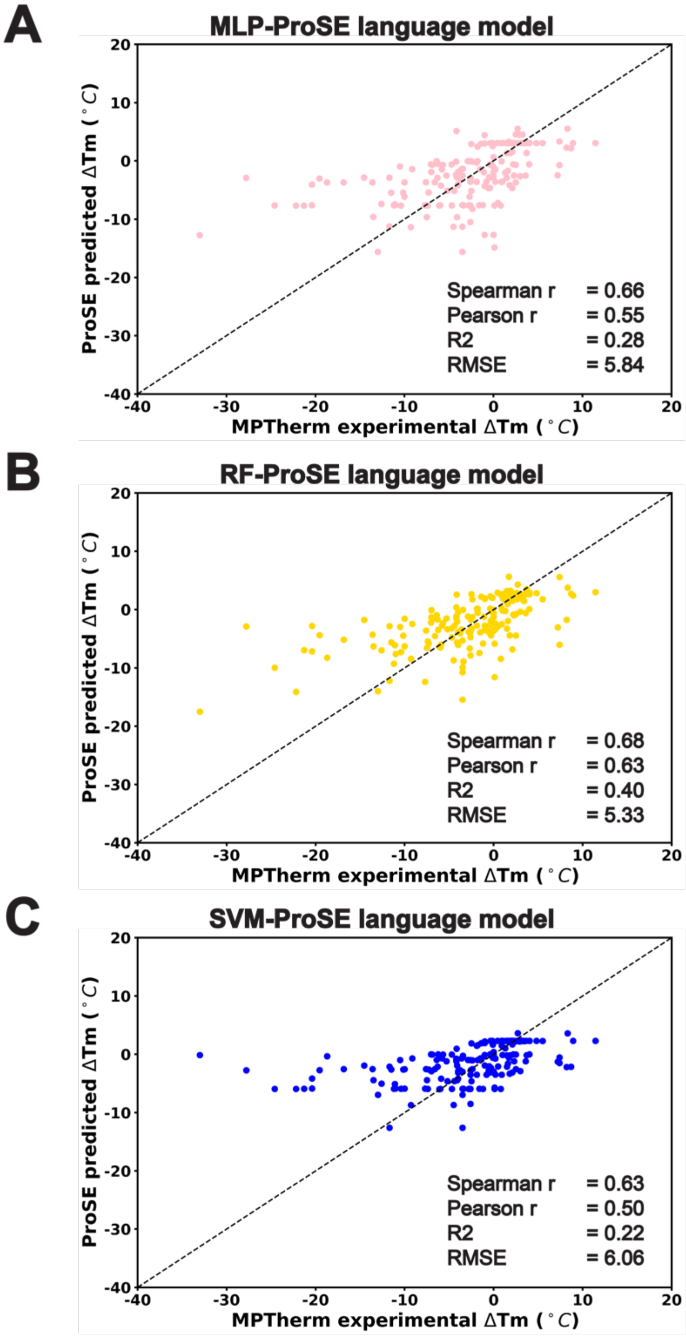


**Figure S8. Performance of different regression models on the MPTherm training dataset (n=917).** The correlation between the predicted (y-axis) and curated experimental $\Delta T_m$ (°C) data (x-axis) for (**a**) MLP, (**b**) RF, and (**c**) SVM. Regression model performance is demonstrated as a scatterplot where the dashed black line refers to a linear correlation of y=x. Statistical metrics for evaluating respective model performance are included within each panel.

**Table S1. Hyperparameter search for SVM, RF and MLP models trained on ProSE embeddings for the MPTherm dataset.**

| Regressor | Hyperparameters Tested | Selected hyperparameter |
|---|---|---|
| SVM | C: [0.1, 1.0, 10.0]<br>kernel: [‘linear’, ‘poly’, ‘rbf’, ‘sigmoid’] | C: 10.0<br>kernel: ‘rbf’ |
| RF | # of estimators: [10, 20, 50, 100]<br>max depth: [50, 100]<br>max features: [‘sqrt’, ‘log2’] | # of estimators: 100<br>max depth: 50<br>max features: ‘log2’ |
| MLP | hidden layer: [(512), (512, 256), (512, 256, 128), (512, 256, 128,64)]<br>batch size: [32, 64]<br>learning rate: [0.01, 0.001] | hidden layer: (512, 256, 128)<br>batch size: 64<br>learning rate: 0.01 |

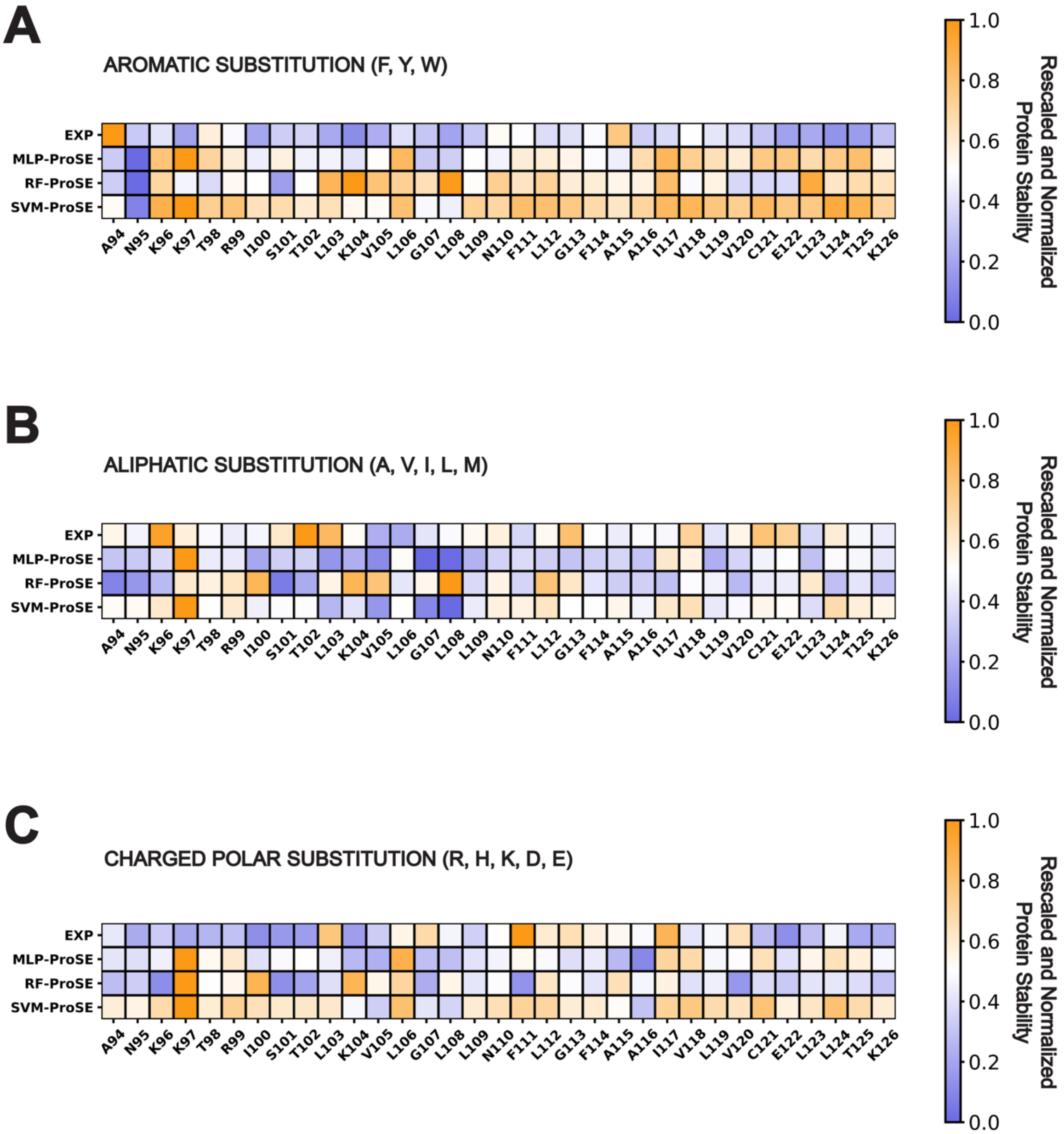


**Figure S9. Evaluating performance of ProSE transfer learning models against AtSWEET13-TM4 DMS data – Part 1.** Rescaled and normalized protein stability values incurred upon mutation are shown for experimental values (EXP) versus *in silico* predictions. The unit scale is expressed from 0 to 1. A value of 0 (dark blue) indicates enhanced predicted thermostability/experimentally reported expression values in response to mutation. A value of 1 (orange) indicates depleted predicted thermostability/experimentally reported expression values in response to mutation. Grey boxes with a red diagonal line pattern indicate that the *in silico* software failed to report a predicted mutation score for all of the possible amino acids enumerated in the substitution category. **a**, Aromatic amino acid substitutions (F, Y, W)**. b,** Aliphatic amino acid substitutions (A, V, I, L, M). **c,** Charged polar amino acid substitutions (R, H, K, D, E).

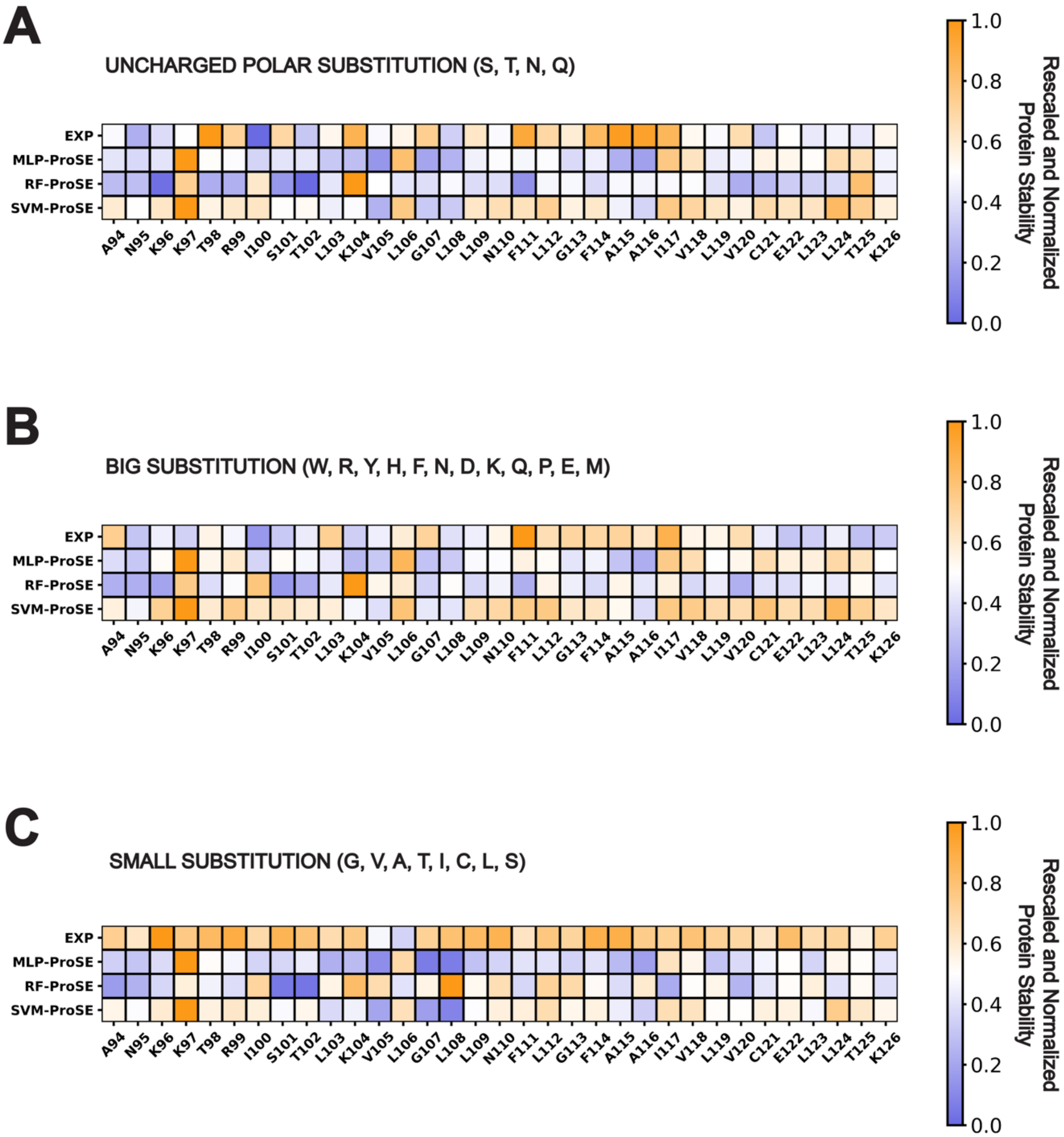


**Figure S10. Evaluating performance of ProSE transfer learning models against AtSWEET13-TM4 DMS data – Part 2.** Rescaled and normalized protein stability values incurred upon mutation are shown for experimental values (EXP) versus *in silico* predictions. The unit scale is expressed from 0 to 1. A value of 0 (dark blue) indicates enhanced predicted thermostability/experimentally reported expression values in response to mutation. A value of 1 (orange) indicates depleted predicted thermostability/experimentally reported expression values in response to mutation. Grey boxes with a red diagonal line pattern indicate that the *in silico* software failed to report a predicted mutation score for all of the possible amino acids enumerated in the substitution category. **a**, Uncharged polar amino acid substitutions (S, T, N, Q)**. b,** Big amino acid substitutions (W, R, Y, H, F, N, D, K, Q, P, E, M). **c,** Small amino acid substitutions (G, V, A, T, I, C, L, S).

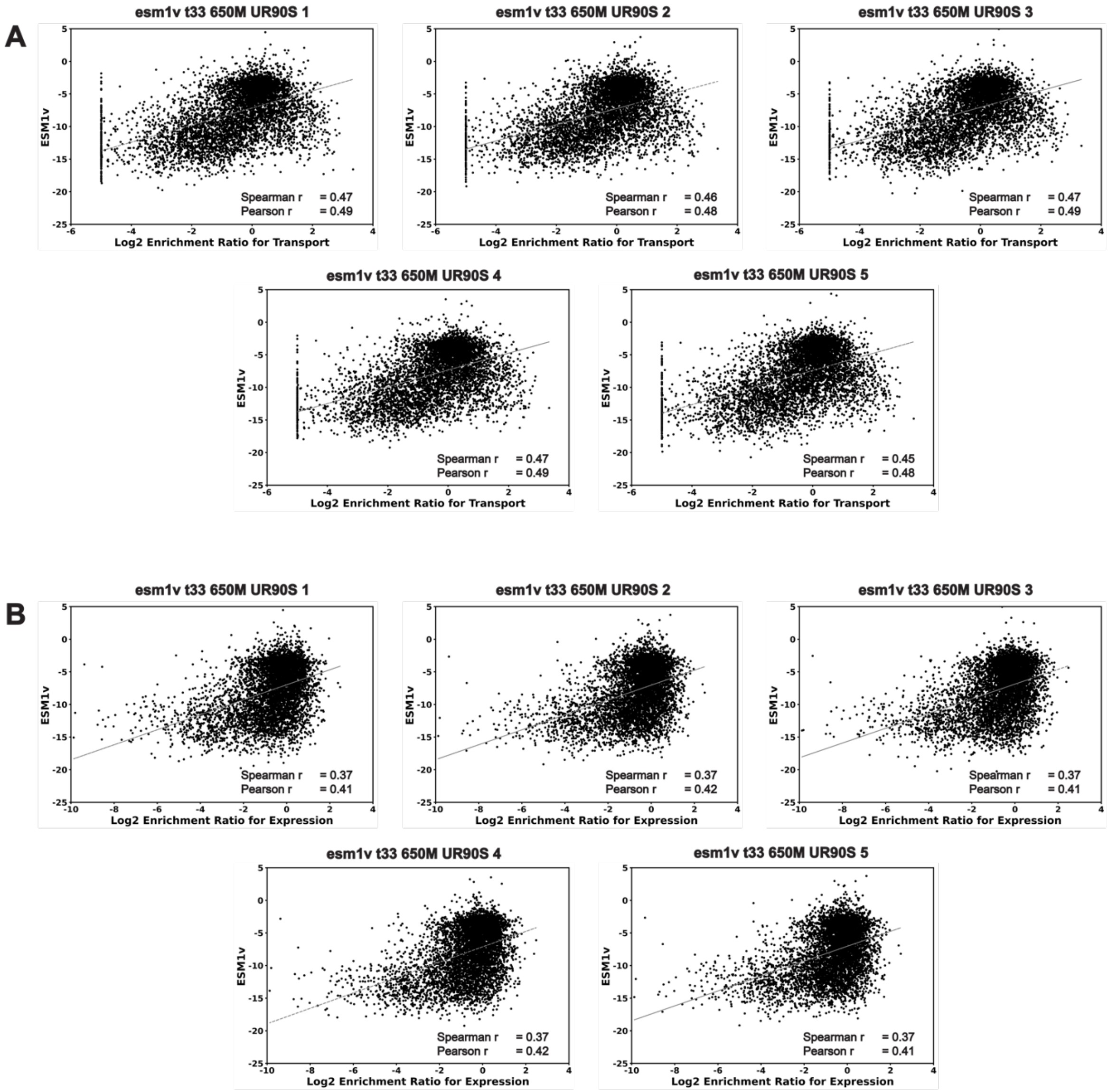


**Figure S11. ESM-1v zero shot learning is unable to accurately reproduce AtSWEET13 DMS data.** Performance for five different pretrained base models of the ESM-1v algorithm are measured against (**a**) $Log_2$ enrichment ratio for AtSWEET13 transport of esculin, and (**b**) $Log_2$ enrichment ratio for AtSWEET13 expression.
